# MAP65-1/PRC1 reinforces microtubules through nucleation

**DOI:** 10.1101/2025.09.14.676082

**Authors:** Mariana Romeiro Motta, Constantin Grisam, Mona Grünewald, Belinda König, Laura Meißner, Lukas Niese, Stefan Diez, Olivier Hamant, Laura Schaedel

**Author notes:** Molecular Biophysics and Biochemistry Department, Yale University, 06511 New Haven, USA.

## Abstract

Microtubules are dynamic cytoskeletal filaments that help shape cells and guide their response to external cues, including mechanical stress. How cells reorganize and reinforce microtubule arrays under stress remains poorly understood. Here, we show that MAP65-1/PRC1 promotes microtubule nucleation on pre-existing lattices *in vitro*, particularly on microtubules with lattice defects and damage sites associated with mechanical stress. MAP65-1 preferentially binds bent microtubules both *in vitro* and in cells, likely because bending induces lattice damage, enhancing nucleation on and bundling of these microtubules *in vitro*. This creates a potential feedback loop where mechanical stress promotes MAP65-1 binding, which in turn stabilizes microtubules by nucleation and bundling and reinforces the alignment of the array with mechanical stress. Thus, we reveal a previously unknown role for MAP65/PRC1 proteins in lattice-based nucleation and suggest a mechanism by which cells record and respond to mechanical stress through microtubule reorganization.

## Introduction

Microtubules form highly organized arrays that play critical roles in intracellular transport, cell division, and morphogenesis. Their array architecture is dynamically sculpted through growth and disassembly^1,2^, severing^3^, cross-linking^4^, and local nucleation^5^. These mechanisms allow cells to build microtubule arrays adapted to their shape and function, and in response to external cues.

Microtubule arrays can reorganize in response to mechanical stress^6^, enabling cells to adapt to external forces. In animal cells, mechanical cues influence microtubule orientation during processes such as epithelial remodeling, cell migration, cell division, or tissue folding^7–10^. In plants, microtubules align with the main tensile stress direction at the cell cortex^6,11,12^, guiding the deposition of cellulose fibers and influencing tissue morphogenesis^13^. Such observations suggest that microtubules act as mechanosensors, relaying external cues into cytoskeletal organization.

While centrosomes and the γ-tubulin ring complex were long considered the primary sites of microtubule nucleation, many differentiated cells – including plant cells^14^, neurons^15^, and epithelial cells^16^ – organize microtubules without centrosomes. Alternative nucleation pathways are therefore critical for building functional microtubule arrays in these cells. Several microtubule-associated proteins (MAPs) have been shown to reduce the critical tubulin concentration for nucleation in solution *in vitro*, suggesting a potential role in the generation of new microtubules^17^. However, whether such nucleation occurs under physiological conditions, and how it is spatially regulated within cells, remains unclear. Existing microtubule lattices may serve as platforms for microtubule nucleation, as observed in structures such as the spindle midzone^18^ or cortical microtubule arrays in plants^19^.

Critical to the formation of both the spindle midzone and plant cortical microtubule array is the MAP65/PRC1 protein family^4,20,21^, which is best known for cross-linking antiparallel microtubules. Beyond cross-linking, the MAP65 family has been shown to increase microtubule flexibility^22^. However, the role of MAP65 proteins in mechanosensitive responses has not been explored.

Here, we show that MAP65-1/PRC1 promote microtubule nucleation on existing microtubule lattices. This activity is enhanced at sites of structural irregularity in the lattice, such as defects or damage sites. These findings suggest a previously unrecognized role of MAP65-1/PRC1 in reinforcing microtubule arrays through lattice-based nucleation. We propose a model in which microtubule alignment with mechanical stress in plant cells generates lattice defects and damage that promote the recruitment of MAP65-1. Once recruited, MAP65-1 nucleates, bundles, and softens microtubules, thereby supporting the alignment of the microtubule array with the principal direction of tensile stress.

## Results

### The MAP65 family promotes microtubule nucleation on existing microtubule lattices

*In vitro* microtubule reconstitution assays offer a controlled and simplified environment to study microtubule dynamics and protein interactions. For that reason, we examined the effect of MAP65-1, a well-known microtubule bundler (Fig. S1a-c), on microtubule dynamics *in vitro*. In control conditions without MAP65-1, polymerized microtubules incubated with free tubulin at concentrations above the polymerization threshold – but well below the threshold for nucleation in solution – exhibit growth only from their free ends (Fig. 1a-c and Fig. S2a). Because we used a low tubulin concentration of 4 µM, microtubule tips grow slowly (Fig. 1c, control). However, in presence of 100 nM MAP65-1, new microtubules appeared along the existing microtubule lattices (Fig. 1c-e). These new microtubules exhibited dynamic instability, growing completely along the “template” microtubules (likely bundled by MAP65-1), and their fluorescence intensity was comparable to the growing microtubule tips (Fig. 1d and e). Increasing the MAP65-1 concentration from 100 to 200 nM led to a significant decrease in the distance between microtubule appearance events, which represents the inverse of spatial frequency (Fig. S2b and c). Hence, we concluded that MAP65-1 can induce microtubule appearance in a concentration-dependent manner.

**Fig. 1:**
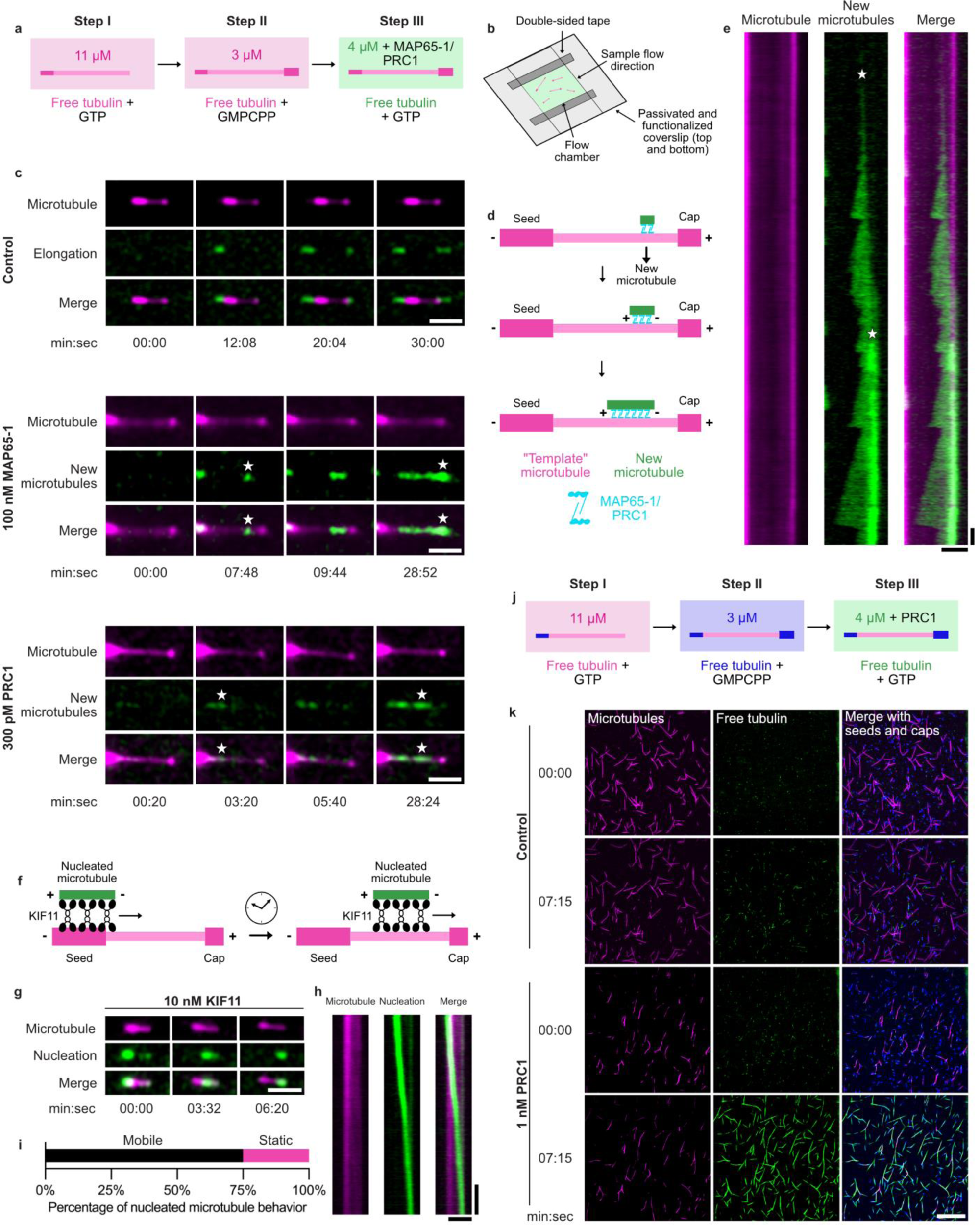
MAP65-1/PRC1 promote microtubule nucleation on existing lattices. **a,** Schematic representation of the experimental setup used to study the function of MAP65-1 and PRC1 in lattice-templated microtubule appearance. Microtubules were grown with ATTO-565-labeled tubulin at a concentration of 11 μM (step I) before they were capped with GMPCPP (step II) and exposed to ATTO-488-labeled free tubulin at a concentration of 4 μM (step III). Step III was observed live for a total of 30 minutes. **b,** Schematic representation of the flow chambers used as an experimental setup throughout the manuscript. 20-μl flow chambers were built from passivated coverslips cut with a diamond pen and attached by double-sided tape. Solutions were sequentially flushed in through one side of the flow chamber. After step III shown in **a**, flow chambers were sealed to prevent evaporation and observed for 30 min. **c,** Microtubule dynamics imaged by TIRFM in the presence of ATTO-488-labeled free tubulin (green) and ATTO-565 microtubules (magenta). In the control (top), only growth from the free microtubule ends (elongation) was observed. In the presence of both 100 nM MAP65-1 (middle) and 300 pM PRC1 (bottom), microtubules appeared and grew in parallel to the existing lattices (microtubule appearance events are indicated with a white star). Scale bars, 5 µm. **d,** Schematic representation of lattice-templated microtubule appearance. Microtubule appearance and growth could be clearly distinguished from elongation at the free ends because it occurred overlapping with the original microtubule lattice “template”. **e,** Kymograph of the corresponding stills in **c** in the presence of MAP65-1. Appearance events are indicated with a white star. Scale bars, 5 µm (horizontal) and 1 min (vertical). **f,** Experimental setup used to test if microtubules that appear in the presence of MAP65-1 are independent from the “template” microtubule lattice. 10 nM KIF11 was added in the presence of 2 μM free tubulin, and some of the new microtubules were effectively used as cargo. **g,** KIF11-mediated microtubule sliding observed by TIRFM after incubation of microtubules with 100 nM MAP65-1 and 4 μM ATTO-488-labeled tubulin. Scale bar, 5 μm. **h,** Kymograph of the corresponding microtubule shown in **g**. Scale bars, 5 μm (horizontal) and 1 min (vertical). **i,** Quantification of nucleated microtubule behavior (mobile or static) upon addition of KIF11 (n = 24 nucleated microtubules, N = 3). **j,** Schematic representation of the experimental setup used to study PRC1-mediated microtubule nucleation in **k.** The same steps were taken as in **a,** except microtubule seeds and caps were polymerized with Alexa-Fluor-647-labeled tubulin (blue) instead of ATTO 565. **k,** Microtubule dynamics in the absence (control) and in the presence of 1 nM PRC1 observed by TIRFM. Scale bar, 50 μm.

Given that the function of MAP65-1 is conserved across the eukaryotic domain, we tested if the human homolog of MAP65-1, PRC1, also promotes microtubule appearance. In the presence of 300 pM PRC1, microtubules appeared and grew on the existing lattices similarly to MAP65-1 (Fig. 1c). Importantly, we worked with MAP concentrations well below those that induce microtubule nucleation in solution. For MAP65-1, we only observed microtubule nucleation in solution at concentrations as high as 500 nM MAP65-1 (Fig. S3); for PRC1, nucleation in solution was detected only above 3 nM PRC1 (Fig. S3). Although MAP65-1 shares only 24% sequence identity with PRC1, it has significantly higher identity with the other Arabidopsis MAP65 family members (Table S1). This suggests that these other MAP65 proteins likely share MAP65-1’s microtubule-generating function.

Previous studies have shown that the MAP SSNA1 promotes microtubule lattice extensions in the form of branches that directly emerge from one or more protofilaments of the mother microtubule^23^. To determine whether the microtubules that appeared in the presence of MAP65-1 and PRC1 were branches of the underlying lattice or appeared following genuine nucleation events, we introduced human kinesin-5 (KIF11) into our assays after a 30-minute incubation of MAP65-1 with microtubules and free tubulin (Fig. 1f-i). Since KIF11 cross-links and slides antiparallel microtubules, the observed mobility of most new microtubules upon KIF11 addition (75%, Fig. 1i) indicates that they were antiparallel and structurally independent of the “template” microtubule. Thus, we confirmed that these microtubules derived from bona fide nucleation events. Since both MAP65-1 and PRC1 preferentially bind to antiparallel microtubules^24^, it is not surprising that most nucleated microtubules are oriented in an antiparallel manner (Fig. 1d).

With a higher PRC1 concentration (1 nM), we observed ubiquitous PRC1-mediated microtubule nucleation and growth that quickly produced thick microtubule bundles (Fig. 1j and k and S4a and b). In the control (no PRC1), after the same timeframe, only short polymerization stretches were observed from the microtubule ends (Fig. 1k). Hence, MAP65-1/PRC1 promote microtubule nucleation on the microtubule lattice.

### MAP65-1 reversibly binds free tubulin when associated with the microtubule lattice

To investigate the mechanism by which MAP65-1 and PRC1 promote microtubule nucleation, we examined whether MAP65-1 can recruit free tubulin to the microtubule lattice. Since microtubule nucleation and growth require GTP, we performed assays in the absence of GTP to prevent polymerization and analyze free tubulin binding (Fig. 2a and S5a and b). Under these conditions, MAP65-1 localized along the microtubule lattice and recruited free tubulin, as hypothesized (Fig. 2b-f).

**Fig. 2:**
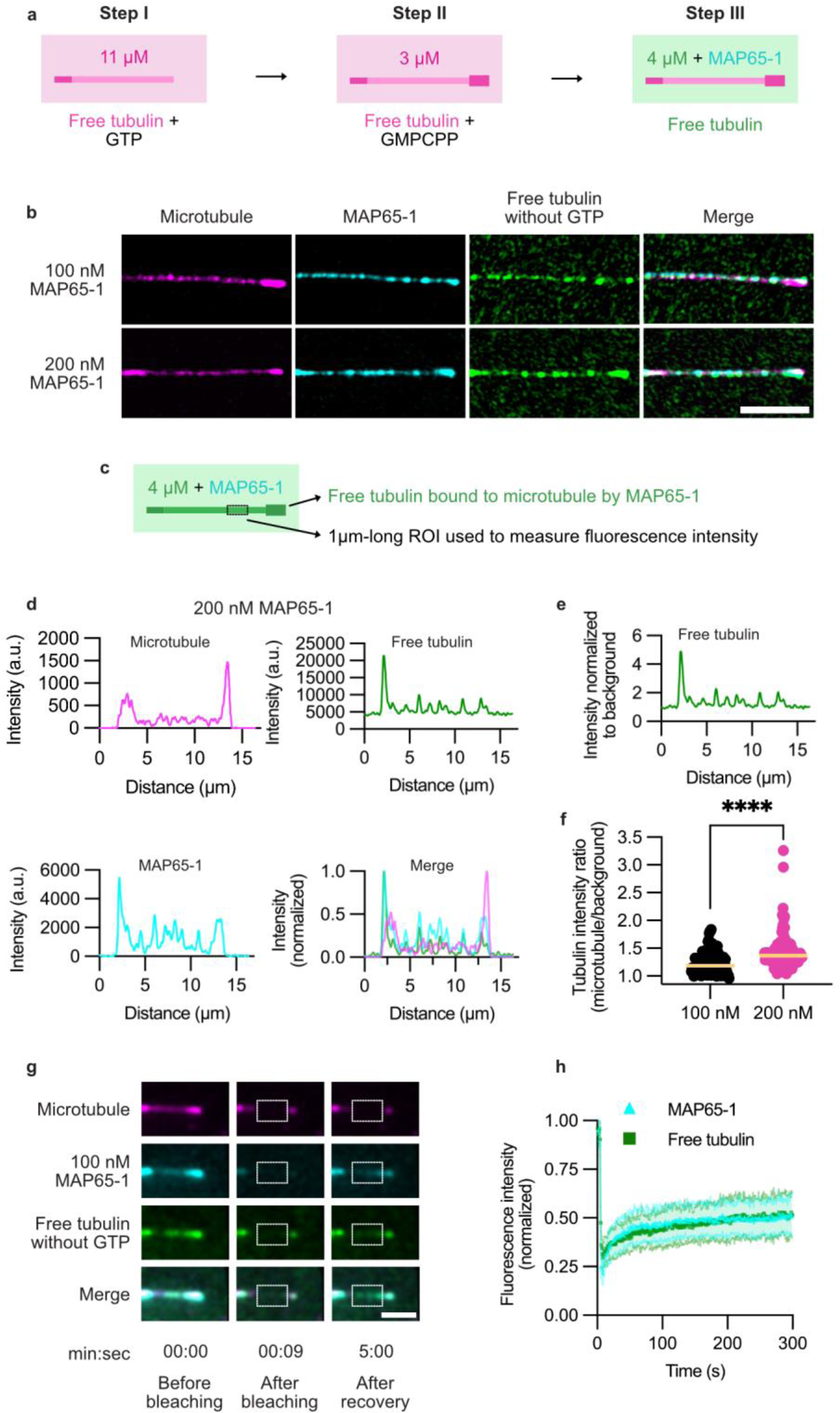
MAP65-1 concentrates tubulin on the microtubule lattice. **a,** Schematic representation of the experimental setup used to study free tubulin recruitment by MAP65-1. Microtubules were grown with Alexa-Fluor-647-labeled tubulin at a concentration of 11 μM (step I) before they were capped with GMPCPP (step II) and exposed to ATTO-565-labeled free tubulin at a concentration of 4 μM and 100 nM or 200 nM GFP-MAP65-1 (step III). **b,** Recruitment of free tubulin in the absence of GTP to the microtubule lattice by 100 and 200 nM GFP-MAP65-1 (cyan) observed by CLSM. Microtubules in magenta and free tubulin in green. Scale bar, 5 μm. **c,** Schematic representation of the setup used to analyze free tubulin recruitment by MAP65-1 to the microtubule lattice. ROIs with a length of 1 µm were drawn to measure fluorescence intensity across the full width of the microtubule, which was then divided to the fluorescence intensity in the background to estimate tubulin concentration. **d,** Graphs represent line scans along the microtubule shown in **b** in the presence of 200 nM GFP-MAP65-1. **e,** Line scan of free tubulin with the intensity normalized to the background free tubulin level. The graph corresponds to the microtubule shown in **b** with 200 nM GFP-MAP65-1 and the corresponding graph in **d**. **f,** Quantification of tubulin recruitment to the microtubule lattice (normalized to the background level) in the presence of 100 nM (n = 150 microtubule segments of 1 µm) and 200 nM (n = 113 microtubule segments of 1 µm) MAP65-1. Bars represent the median values. Statistics: Mann-Whitney test, **** P < 0.0001. **g,** FRAP assay observed by TIRFM. The dashed white rectangle indicates the bleached region where fluorescence intensity was measured over time. Scale bar, 5 μm. **h,** Graph with icons representing the average normalized fluorescence intensity recovery over time for both GFP-MAP65-1 (cyan) and free tubulin (green; n = 17 microtubules). Shaded region above and below the icons corresponds to the standard deviation for each time point.

Quantitative analysis revealed that, at 200 nM MAP65-1, the average enrichment of free tubulin compared to the solution was approximately 43% (Fig. 2f). However, in our system, a ∼3.5-fold enrichment in tubulin concentration is typically required to initiate nucleation in solution in the absence of MAPs. Although we occasionally observed localized patches where tubulin enrichment approached this threshold, such events were considerably less frequent than the nucleation events observed, as reflected by the measured distance between nucleation events under the same MAP65-1 concentration (Fig. S2b and c). One possibility is that small nucleation intermediates fall well below our resolution limit, leading us to underestimate local tubulin concentration. Yet, we cannot exclude that mechanisms other than tubulin concentration may play a role in the observed microtubule nucleation.

To determine whether MAP65-1 and free tubulin bound to the microtubule lattice are dynamically exchanged with molecules in solution, we performed fluorescence recovery after photobleaching (FRAP) experiments in the absence of GTP. Following photobleaching, both GFP-MAP65-1 and free tubulin signals partially recovered on the microtubule lattice (within 300 s; Fig. 2g and h), indicating that both proteins undergo dynamic exchange with the soluble pool.

### MAP65-1 recognizes bent microtubules in cells and *in vitro*

Our *in vitro* data demonstrate that MAP65-1 promotes microtubule nucleation on existing lattices, at least partly by recruiting free tubulin that undergoes exchange with the soluble protein pool. Yet, the relevance of this activity and its spatial regulation in cells remained unclear.

Because microtubules are stiff polymers, they would tend to align with the lowest curvature cell axis by default. This is observed in non-pressurized wall-less plant cells^25^ or in non-growing hypocotyl cells^26^. However, in turgid and growing plant cells, cortical microtubules often align with the highest curvature axis instead. Because the highest curvature axis for a pressurized cylindrical cell also corresponds to the maximal tensile stress direction, it was proposed that tensile stress overrides geometrical cues^25^. Since MAP65-1 makes microtubules softer, we reasoned that microtubules would be less responsive to the lowest curvature cell axis and more responsive to mechanical stress in the presence of MAP65-1. Thus, we wondered whether MAP65-1 can specifically promote microtubule nucleation on existing, bent microtubules, leading to a reinforcement of this specific microtubule subset. To investigate this, we observed the distribution of MAP65-1 in cells of young *Arabidopsis* stems containing a microtubule (GFP-MBD) and a MAP65-1 (MAP65-1-mCherry) reporter. MAP65-1 fluorescence intensity was around twice as high on bent microtubules in comparison to straight ones (Fig. 3a-c and Table S2), and we observed that both MAP65-1 and MBD fluorescence intensity positively correlated with maximum microtubule curvature (Fig. 3d and e). Note that it remains unclear whether MAP65-1 binds to pre-existing bundles that are more prominent in highly curved regions, or whether it contributes to bundle formation in these regions by favoring microtubule nucleation and bundling in highly bent microtubules.

**Fig. 3:**
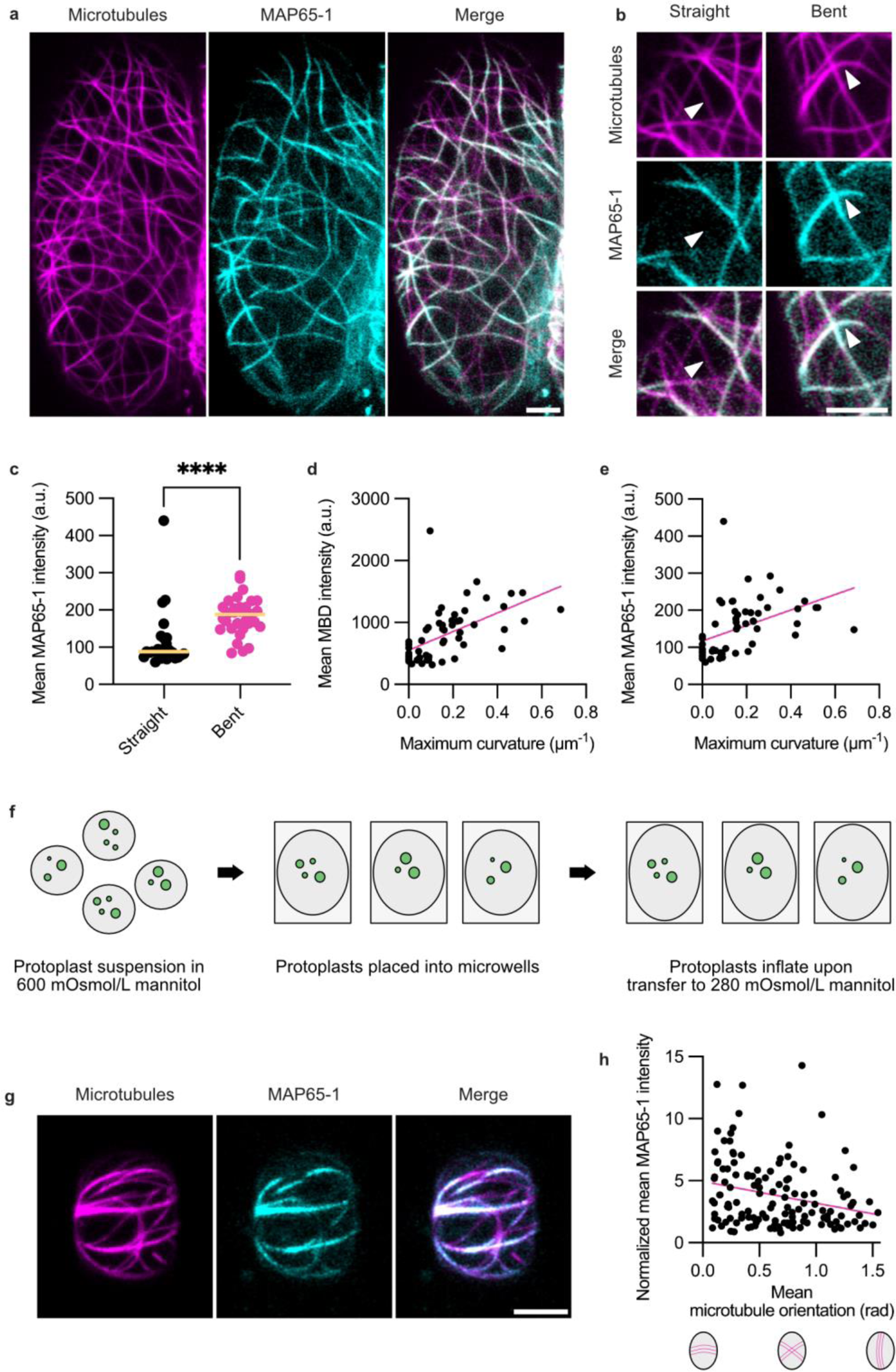
MAP65-1 preferentially localizes to bent microtubules in cells. **a,** Average intensity Z-projection of CLSM of epidermal hypocotyl cells from 5-day-old plants co-expressing *p35S::GFP-MBD* and *pMAP65-1::MAP65-1-mCherry*. Scale bar, 5 μm. **b,** Arrowheads indicate examples of straight and bent microtubules extracted from the image shown in **a.** Scale bar, 5 μm. **c,** Quantification of mean MAP65-1 intensity on straight (maximum curvature < 0.1 μm^-1^, n = 27) and bent (maximum curvature > 0.1 μm^-1^, n = 31) microtubules, N = 5 cells. Bars indicate the median values. Statistics: Mann-Whitney test, **** P < 0.0001. **d,** Quantification of mean MBD intensity per microtubule against microtubule maximum curvature. The pink line indicates a simple linear regression (Spearman r = 0.67, P < 0.0001, n = 58 microtubules). **e,** Quantification of mean MAP65-1 intensity per microtubule against microtubule maximum curvature. The pink line indicates a simple linear regression (Spearman r = 0.64, P < 0.0001, n = 58 microtubules). **f,** Schematic representation of the protoplast experiment. Protoplasts are first extracted in a solution with 600 mOsmol**/**L mannitol, then allowed to sediment into rectangular microwells followed by an exchange to a solution with 280 mOsmol/L mannitol that causes the protoplasts to inflate. **g,** Average intensity Z-projection of CLSM of enclosed protoplasts extracted from roots of plants co-expressing *p35S::GFP-MBD* and *pMAP65-1::MAP65-1-mCherry*. Scale bar, 5 μm. **h,** Quantification of normalized mean MAP65-1 intensity against mean microtubule orientation. The pink line indicates a simple linear regression (Spearman r = -0.25, P = 0.0028, n = 135 microtubules).

The analyses above in plant tissues give an indication of MAP65-1’s ability to recognize bent microtubules. However, due to the complexity of the tissue, including water fluxes and different turgid statuses for instance, these analyses only offer a correlation. To get closer to causality, we used a much simpler system, consisting of wall-less plant cells (protoplasts) confined in rectangular microwells (Fig. 3f). Upon osmotic pressurization, achieved by transferring cells to a medium with lower osmolarity, microtubules have been shown to reorient from their default longitudinal orientation to a transverse alignment, which corresponds to the highest curvature and predicted highest tension axis^25^. Under these conditions, MAP65-1 preferentially accumulated on microtubules aligning transversely (Fig. 3g). This was supported by a significant, albeit weak (Spearman r = -0.25), negative correlation between normalized mean MAP65-1 intensity and mean microtubule orientation two hours after pressurization (Fig. 3h), indicating that MAP65-1 is enriched on mechanically challenged microtubules. Microtubules may thus be able to disentangle geometry from mechanical stress through a feedback loop where MAP65-1 recruitment allows microtubule softening, which, in turn, promotes alignment with tensile stress, bending along the highly curved axis of the cell, and further MAP65-1 recruitment.

In cells, MAP65-1 accumulation on bent microtubules may result from different factors, such as the curvature of the microtubule itself, mechanical stress, or the recruitment of other MAPs. Hence, to dissect these factors, we tested in our *in vitro* assays if MAP65-1 could recognize microtubule curvature alone. To do that, we bent microtubules by using fluid flow and maintained them in a bent conformation by attaching the microtubule cap to the coverslip. Next, we incubated microtubules with GFP-MAP65-1. We found that MAP65-1 accumulated around 3 times more on bent microtubules in comparison to straight ones (Fig. 4a and b). Accordingly, mean MAP65-1 intensity significantly correlated with maximum microtubule curvature (Fig. 4c).

**Fig. 4:**
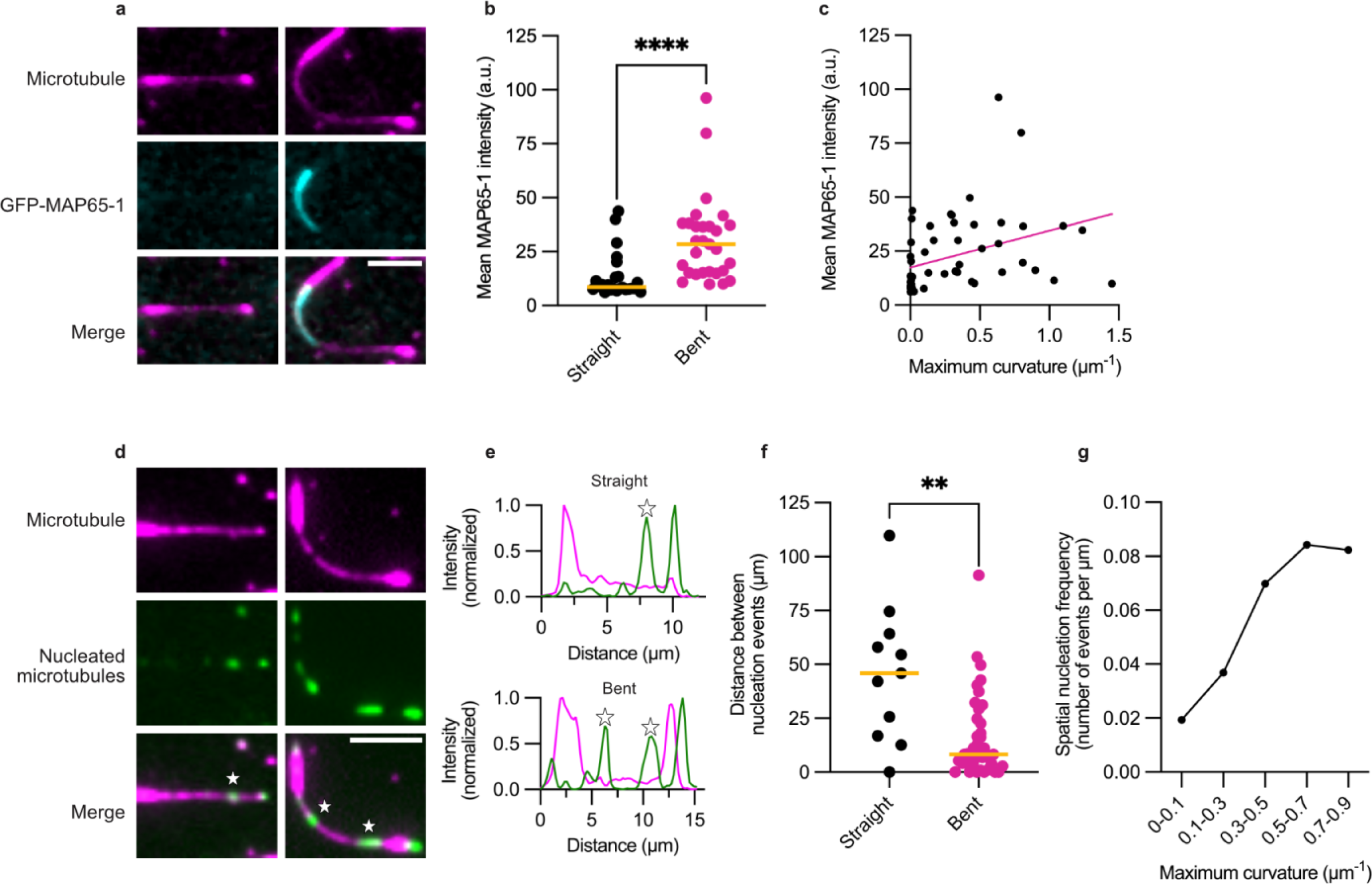
MAP65-1 promotes microtubule nucleation on bent microtubules *in vitro*. **a,** Microtubules attached to the surface at both ends, straight or forced to statically bend by fluid flow, observed by TIRFM. 100 nM MAP65-1 was added at a ratio of 15% GFP-MAP65-1 to 85% non-fluorescent MAP65-1. Scale bar, 5 μm. **b,** Quantification of mean MAP65-1 intensity on straight (maximum curvature < 0.1 μm^-1^, n = 24) and bent (maximum curvature > 0.1 μm^-1^, n = 29) microtubules, N = 3. Bars indicate the median values. Statistics: Mann-Whitney test, **** P < 0.0001. **c,** Quantification of mean MAP65-1 intensity per microtubule against microtubule maximum curvature. The pink line indicates a simple linear regression (Spearman r = 0.52, P < 0.0001, n = 53). **d,** Microtubule nucleation mediated by 100 nM MAP65-1 on straight and bent microtubules observed by TIRFM. Nucleation events are indicated with a white star. Scale bar, 5 μm. **e,** Graphs represent line scans along the microtubules shown in **d** (original microtubule in magenta, nucleated/elongated microtubules in green). **f,** Quantification of the distance between nucleation events on straight (maximum curvature < 0.1 μm^-1^, n = 11 events) and bent (maximum curvature > 0.1 μm^-1^, n = 40 events) microtubules. Bars indicate the median values. Total microtubule lengths of 723.52 μm (straight) and 819.45 μm (bent) were analyzed, N = 3. Statistics: Mann-Whitney test, ** P = 0.0012. **g,** Spatial nucleation frequency (number of events per µm) according to microtubule maximum curvature (n = 14 events for 0–0.1 µm^-1^, 17 events for 0.1–0.3 µm^-1^, 10 events for 0.3–0.5 µm^-1^, 8 events for 0.5–0.7 µm^-1^ and 6 events for 0.7–0.9 µm^-1^). Total microtubule lengths = 723.52 µm (0–0.1 µm^-1^), 461.18 µm (0.1–0.3 µm^-1^), 143.07 µm (0.3–0.5 µm^-1^), 94.91 µm (0.5–0.7 µm^-1^) and 72.87 µm (0.7–0.9 µm^-1^).

Next, we tested if MAP65-1-mediated microtubule nucleation happened more frequently on bent microtubules as well. We incubated microtubules with free tubulin and 100 nM MAP65-1 for 30 min followed by washing free tubulin away and microtubule stabilization with taxol (Fig. 4d-g). The distance between nucleation events was approximately 5 times smaller in bent microtubules in comparison to straight microtubules (Fig. 4f). Average spatial nucleation frequency also increased with maximum microtubule curvature (Fig. 4g). Notably, microtubules nucleated also in regions other than those of highest curvature along bent microtubules (Fig. S6), which suggests that additional factors likely influence the site of nucleation.

### MAP65-1 does not recognize microtubule curvature through lattice expansion or compaction

MAPs that recognize microtubule curvature often exhibit preferential binding to either expanded or compacted lattice conformations because a curved microtubule has an expanded outer surface and a compacted inner surface^27,28^.

Interestingly, GMPCPP-polymerized microtubules, which adopt an expanded lattice conformation^29^, were the predominant sites of nucleation in the presence of MAP65-1, despite comprising only a small fraction of the microtubule length (Fig. S7a and b). To test whether MAP65-1 preferentially binds to GMPCPP-containing microtubules, we quantified GFP-MAP65-1 intensity on GMPCPP seeds versus GDP lattice regions. We observed a significant enrichment of MAP65-1 on the seeds (Fig. S7c-e), indicating a preference for the GMPCPP lattice.

To determine whether MAP65-1 specifically recognizes the expanded lattice state, we treated microtubules with taxol, which also induces lattice expansion^29^. However, taxol-treated microtubules showed significantly reduced MAP65-1 accumulation (Fig. S7f and g), suggesting that lattice expansion alone does not account for MAP65-1 binding.

Since some MAPs that recognize specific lattice states can also induce lattice expansion or compaction upon binding^30^, we examined microtubules incubated with GFP-MAP65-1 using microfluidics (Fig. S7h and i). Even at a high concentration (500 nM), MAP65-1 did not induce measurable changes in lattice length. Thus, we concluded that MAP65-1 does not recognize microtubule curvature through lattice expansion or compaction.

### MAP65-1 and PRC1 recognize microtubules with structural irregularities

In our experiments, we define lattice defects as irregularities that arise during polymerization (for example, a change in protofilament number), whereas damage refers to alterations that occur after polymerization, such as local tubulin loss or breaks in the lattice. Bent microtubules are particularly prone to lattice damage, as reflected by increased tubulin turnover along their shafts^31^.

Given MAP65-1’s ability to recognize bent microtubules, we thus hypothesized that MAP65-1/PRC1 might detect structural irregularities in the microtubule lattice, such as defects and damage. To test this, we manipulated the frequency of microtubule irregularities using two approaches: (1) by reducing the tubulin concentration for polymerization to half of that used in fast-growth conditions, generating microtubules with fewer defects^32^ (referred to as slow growth conditions); and (2), by treating microtubules with taxol to induce lattice damage^33^, followed by thorough washing to remove the drug (Fig. 5a-g).

**Fig. 5:**
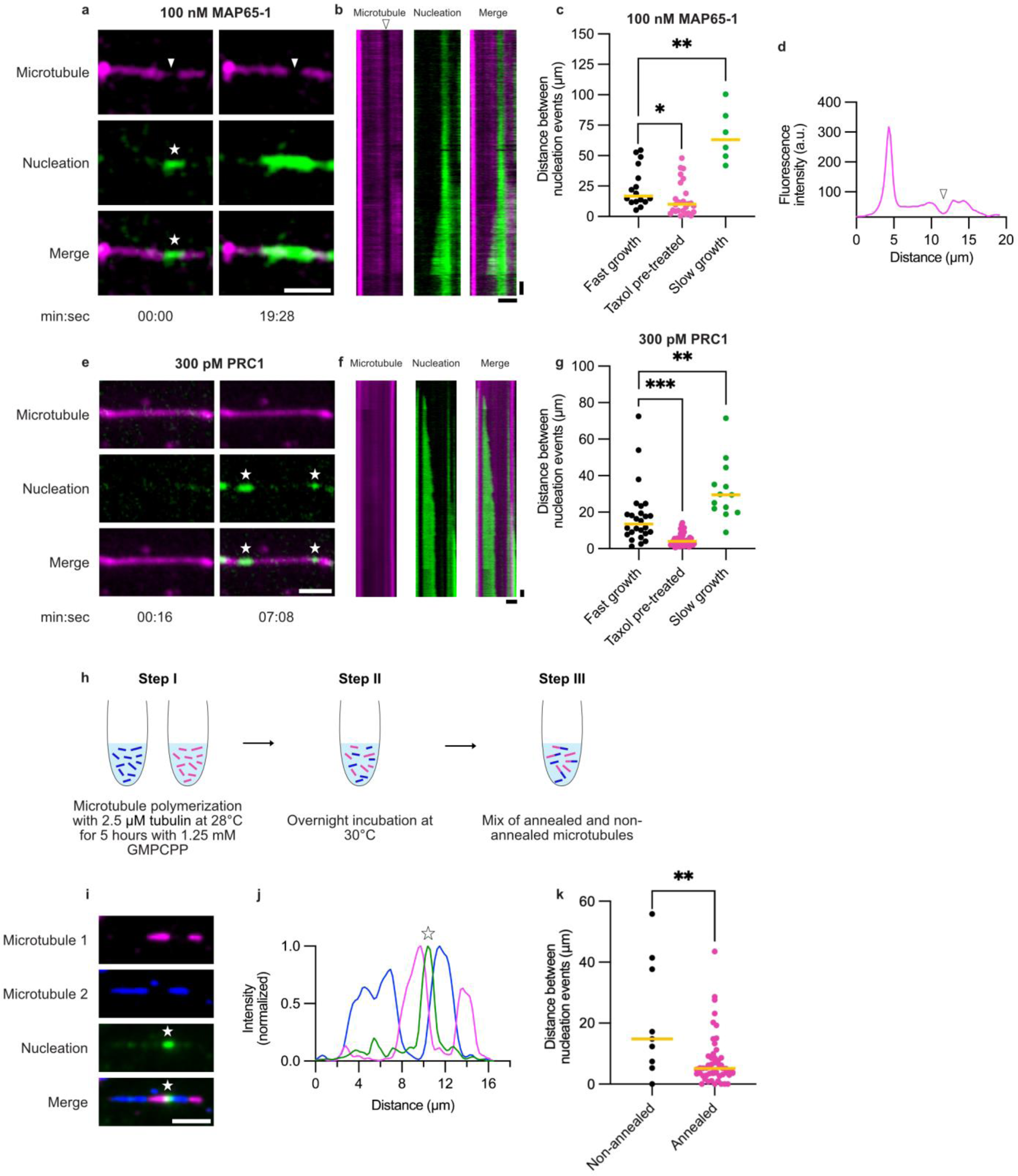
MAP65-mediated microtubule nucleation depends on the amount of microtubule defects. **a,** Microtubule nucleation on taxol pre-treated microtubule mediated by 100 nM MAP65-1 imaged by TIRFM. Free tubulin (green) labeled with ATTO 488 and microtubules (magenta) with ATTO 565. White arrowhead indicates presumed microtubule damage site from where nucleation (white star) happens. Scale bar, 5 µm. **b,** Kymograph of the corresponding microtubule shown in **a**. Scale bars, 5 µm (horizontal) and 1 min (vertical). **c,** Quantification of the distance between nucleation events on fast grown, taxol pre-treated and slowly grown microtubules. A total of 16 (fast growth), 26 (taxol pre-treated) and 6 (slow growth) nucleation events were observed. Bars indicate the median values. Total microtubule lengths of 430.04 µm, 398.67 µm and 453.21 µm were analyzed respectively, N ≥ 3. Statistics: Mann-Whitney test, Bonferroni-corrected, * P = 0.015 and ** P = 0.0016. **d,** Graph represents line scan along the microtubule shown in **a**, showing fluorescence intensity at presumed damage site (white arrowhead). **e,** Microtubule nucleation on taxol pre-treated microtubule mediated by 300 pM PRC1 imaged by TIRFM. Free tubulin (green) labeled with ATTO 488 and microtubules (magenta) with ATTO 565. Nucleation events are indicated with white stars. Scale bar, 5 µm. **f,** Kymograph of corresponding microtubule shown in **e**. Scale bars, 5 µm (horizontal) and 1 min (vertical). **g,** Quantification of the distance between nucleation events on fast grown, taxol pre-treated and slowly grown microtubules. A total of 26 (fast growth), 46 (taxol pre-treated) and 13 (slow growth) nucleation events were observed. Bars indicate the median values. Total microtubule lengths of 536.38 µm, 254.24 µm and 446.95 µm were analyzed respectively, N ≥ 3. Statistics: Mann-Whitney test, Bonferroni-corrected, ** P = 0.0024 and *** P = 0.0002. **h,** Schematic illustration of the experimental setup to generate annealed microtubules. Microtubules were slowly polymerized with GMPCPP and in the presence of ATTO-565-labeled or Alexa-Fluor-647-labeled tubulin (step I). The two microtubule populations were then mixed and incubated overnight at 30°C (step II). The next day, the mix of non-annealed and annealed microtubules was used for the experiments (step III). **i,** Microtubule nucleation mediated by 100 nM MAP65-1 observed on annealed microtubules by TIRFM. Nucleation event is indicated with white star. Scale bar, 5 µm. **j,** Graph represents line scan along the microtubule shown in **i** (microtubule 1 in magenta, microtubule 2 in blue, nucleated microtubule in green). **k,** Quantification of the distance between nucleation events on non-annealed and annealed microtubules. A total of 9 (non-annealed) and 54 (annealed) nucleation events were observed. Bars indicate the median values. Total microtubule lengths of 244.35 µm and 406.52 µm were analyzed respectively, N = 2. Statistics: Mann-Whitney test, ** P = 0.0099.

In taxol pre-treated microtubules, we occasionally observed regions with noticeably reduced lattice fluorescence (Fig. 5a-d, white arrowhead), which likely represent damage sites. This observation is consistent with previous reports showing that treatment with taxol overnight at room temperature causes considerable damage to the microtubule lattice^33^. Notably, MAP65-1-mediated microtubule nucleation events often originated from these damage sites. The median distance between nucleation events was significantly reduced in taxol-pre-treated microtubules (10.06 µm) compared to fast-growth conditions (16.88 µm; P = 0.015, Fig. 5c). In contrast, under slow-growth conditions, the median distance between nucleation events increased to 63.00 µm (P = 0.0016). This corresponds to a spatial frequency nearly four times lower than that observed in fast-growth conditions.

We validated these findings in an alternative approach, in which GMPCPP-stabilized microtubules were polymerized under high- and low-defect conditions^34^, followed by incubation with MAP65-1 and free tubulin (see Methods, Fig. S8a). Under low-defect conditions, the median distance between nucleation events significantly increased from 3.44 µm to 6.74 µm, nearly a twofold difference, compared to the high-defect regime (Fig. S8b and c). These results indicate that the spatial frequency of MAP65-1-mediated microtubule nucleation is promoted by the presence of structural irregularities in the lattice.

To determine whether this function is conserved, we tested whether PRC1 also promotes nucleation in a defect- and damage-dependent manner. We found that PRC1-mediated nucleation occurred approximately three times more often on taxol-pre-treated microtubules and about two times less often on microtubules grown under slow-growth conditions (Fig. 5e-g). We concluded that both MAP65-1 and PRC1 preferentially promote microtubule nucleation on lattices with increased structural irregularity.

### MAP65-1 mediated microtubule nucleation often co-localizes with annealing sites

A remaining question was whether MAP65-1/PRC1 promote microtubule nucleation directly at defect or damage sites. To test this, we generated microtubules with defined annealing sites, which often form structural defects due to the imperfect alignment of frayed microtubule ends. We polymerized two populations of GMPCPP-stabilized microtubules, mixed them in equal amounts and allowed them to anneal overnight (Fig. 5h). This approach generated a mixed population of annealed and non-annealed microtubules. We then incubated the mixture with MAP65-1 and free tubulin, followed by washing and stabilization with taxol.

Strikingly, approximately 45% of the nucleation events co-localized with annealing sites (25 out of 55 events; Fig. 5i and j). This was significantly higher than expected from randomly distributed nucleation sites: among 55 randomly selected microtubule patches, only 6 contained nucleation events (Fisher’s exact test, P < 0.0001). Furthermore, the median distance between nucleation events significantly decreased in annealed microtubules (5.11 µm) compared to non-annealed ones (14.86 µm; P = 0.0099, Fig. 5k), almost a three-fold difference. Thus, we concluded that nucleation occurs at annealing sites more frequently than expected by chance, supporting the hypothesis that MAP65-1/PRC1 can recognize structural defects, where they promote microtubule nucleation.

### Bundled microtubules are more resistant to breakage after free tubulin removal

Considering that MAP65-1 preferentially promotes nucleation – and thus bundling – on microtubules with structural defects and damage, we wondered whether microtubules bundled by MAP65-1 are more resistant to breakage. To test this, we removed free tubulin from our *in vitro* assays, a condition that causes microtubules to gradually lose tubulin dimers from their lattices, soften, and eventually break. After polymerizing and capping microtubules, we simultaneously removed free tubulin and added 100 nM MAP65-1 and imaged microtubules over time (Fig. 6a and b). Remarkably, bundled microtubules survived almost 3 times longer compared to single ones (Fig. 6b), showing that MAP65-1 protects microtubule bundles from breakage.

**Fig. 6:**
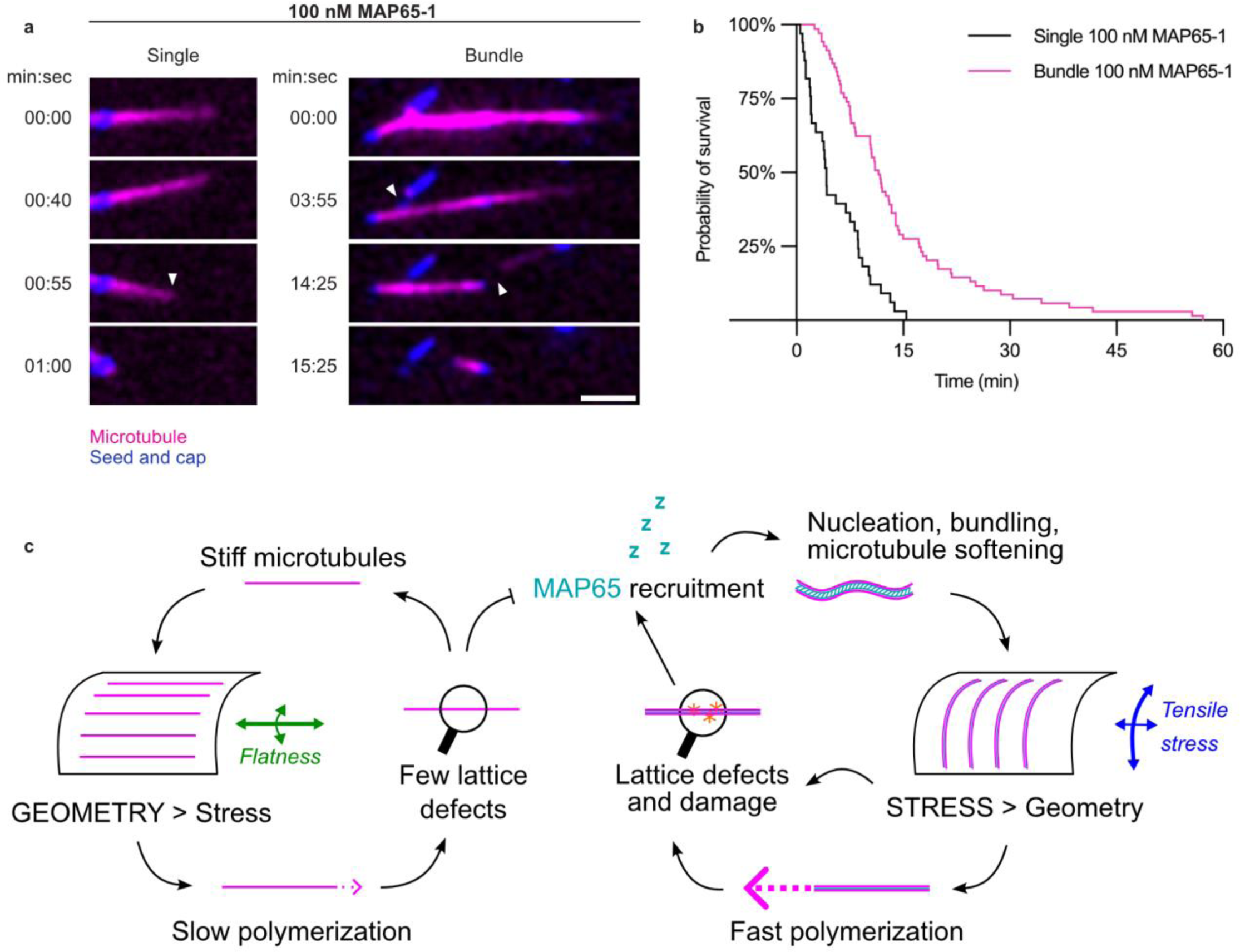
Bundled microtubules survive longer in the absence of free tubulin. **a,** Microtubule dynamics upon removal of free tubulin and incubation with 100 nM MAP65-1 imaged by TIRFM (microtubules labeled with ATTO-565 in magenta, seeds and caps in blue). White arrowheads indicate microtubule breakage. Scale bar, 5 µm. **b,** Quantification of microtubule survival over time upon removal of free tubulin in the presence of 100 nM MAP65-1 for single (N = 8, n = 33 microtubules) and bundled (N = 5, n = 69 microtubules) microtubules. Statistics: Logrank (Mantel-Cox) test, **** P < 0.0001. **c,** Schematic representation of the hypothetical function of MAP65-1 in reinforcing microtubule alignment with mechanical stress. Microtubules polymerize faster under tensile stress (lower right part of the panel); this induces more defects in the lattice (along with more damage sites due to the high curvature of these microtubules); these defects and damage sites recruit MAP65, which then stabilizes, nucleates on and bundles microtubules. MAP65 binding also leads to microtubule softening. The alignment of microtubule arrays is the emerging property of their stiffness: stiff microtubules tend to align along the flattest direction of the cell cortex. For soft microtubules, this emerging property is disrupted, and microtubules instead align to a greater extent along the direction of maximal tensile stress, i.e., the surface experiencing the highest curvature for a pressurized cell. These mechanisms bring forth a positive feedback loop, where then microtubules under stress again polymerize faster, have more defects (and damage due to bending) and recruit MAP65. MAP65 thus protects microtubule under stress and promotes its own recruitment.

Mechanical stress in the form of tension has been shown to accelerate microtubule polymerization *in vitro*^35^. Based on our findings that MAP65-1 promotes nucleation on microtubules with more lattice defects and damage, we propose a model in which mechanical stress enhances microtubule polymerization speed in plant cells. Under higher polymerization speeds, microtubules accumulate structural irregularities, for instance in the form of lattice defects, which are recognized by MAP65-1. Microtubules aligning with mechanical stress in plant cells also likely accumulate lattice damage caused by bending. MAP65-1 thus preferentially stabilizes, softens, nucleates on, and bundles these microtubules. As a consequence, MAP65-1 activity promotes microtubule array alignment along mechanical stress patterns (Fig. 6c).

## Discussion

In summary, we showed that MAP65-1 preferentially binds to and promotes nucleation on bent microtubules, supporting a model in which MAP65-1 acts as a mechanosensitive regulator of cytoskeletal organization. This is particularly relevant in plant cells, where cortical microtubules typically align with tensile stress patterns rather than local geometric cues. For instance, in many plant tissues, adjacent cells with different geometries exhibit consistent supracellular microtubule alignment^11,36^. Our finding provides a scenario for microtubules to be differentially sensitive to geometrical and mechanical cues. In turgid cells, tensile stress would promote fast microtubule polymerization and the formation of defects in the lattice; the subsequent recruitment of MAP65-1 would secure the viability of defective microtubules through microtubule addition (nucleation and bundling). As a consequence of MAP65 recruitment, MAP65-decorated microtubules would also become softer. They would then be less inclined to align with the flattest part of the cell cortex (compared to when they were stiff) and instead align more in the direction of tensile stress at the cell cortex, i.e., under high curvature, in a feedback loop. Importantly, we do not expect single microtubules to reorient and align with the direction of highest tensile stress or curvature; instead, a few microtubules stochastically growing in that direction are expected to be sufficient to support the proposed feedback loop.

The co-localization of nucleation events with microtubule annealing sites supports the idea that MAP65-1 recognizes structurally vulnerable regions and reinforces them through nucleation and bundling. Importantly, we demonstrate that microtubules bundled by MAP65-1 are more resistant to breakage following tubulin depletion, suggesting a stabilizing role for MAP65-1-mediated bundling under stress conditions (Fig. 6c). Interestingly, an alternative scenario where tensile stress would promote the expanded lattice conformation of microtubules and further recruitment of MAP65 was not supported by our experiments. MAP65-1-dependent stress perception would thus be mediated through the accumulation of defects and lattice damage. In this scenario, MAP65-1 recruitment to damaged microtubules would serve a dual purpose: it would acutely reinforce microtubules in response to mechanical stress, and, because the resulting bundles are particularly stable, it would also leave a lasting record of past stress events within the cell. This would, in turn, make the microtubule array align better with stress.

The exact molecular mechanism by which MAP65-1/PRC1 promote microtubule nucleation remains to be elucidated. Since the tubulin concentration on the lattice was not high enough to explain the spatial frequency of nucleation activity, it is likely that MAP65-1/PRC1 stabilize nucleation intermediates by enhancing lateral or longitudinal affinity between tubulin dimers^17^. Accordingly, the yeast homolog Ase1 has been proposed to reduce the detachment of terminal tubulin subunits at depolymerizing microtubule tips^37^, which shows that MAP65-1/PRC1 could influence the tubulin dissociation rate from the microtubule lattice.

Note that further synergies may be envisioned here. In particular, higher osmolarity does not only reduce cortical tension, but also leads to reduced microtubule growth^38^. Thus, in non-pressurized cells, slow-growing microtubules might experience less defects in their lattice, leading to reduced recruitment of MAP65-1. The resulting stiffer microtubules would be more sensitive to cell geometry and align with the flat part of the cell cortex, by default.

The mechanism by which MAP65-1 recognizes bent microtubules seems to differ from other MAPs that have been shown to bind bent microtubules through a preference for the expanded or compacted lattice states^27,28,39^. Although we did not observe preferential binding to the taxol-expanded lattice, MAP65-1 exhibited a preference for the GMPCPP lattice. This suggests that MAP65-1 may recognize bent microtubules through alternative features of the microtubule lattice that are not explained solely by expansion or compaction.^28,37,38^ Furthermore, one interesting possibility is that MAP65-1 detects microtubules under tension, a hypothesis that remains to be tested.

Because MAP65-1/PRC1 have not been described to preferentially bind GTP-tubulin, the recognition of lattice defects also seems different from other MAPs that recognize defect sites through the recognition of GTP-tubulin incorporation, like CLASP, CLIP-170, and the EBs^40–43^. Moreover, MAP65-1 may recognize bent microtubules through their increased lattice damage, as bent microtubules show much higher levels of repair (with an increased incorporation of tubulin from solution) along their lattice than straight ones^31^.

Finally, the interplay of MAP65-1/PRC1 with other MAPs that are responsive to mechanical stress is likely complex in cells. For example, PRC1 and Ase1 interact with and recruit CLASP^44,45^, and microtubule bundles formed by MAP65-1 are protected from severing by KATANIN^46^. Recent simulations of plant cortical microtubule arrays also support a key role for bundling in helping microtubule self-organization together with KATANIN’s severing function^47^. Therefore, MAP65-1 might further contribute to the microtubule response to mechanical stress in cells by recruiting CLASP and counter-acting the severing by KATANIN. Overall, it will be interesting to test how depletion of MAP65 in plant cells can affect their response to changes in mechanical stress, although this is a challenging endeavor because the MAP65 family has nine members in *A. thaliana*^48^. Future work could address whether other MAP65 members also localize to bent microtubules or promote nucleation, clarifying whether they have a conserved or specialized role.

Together, our findings position MAP65-1 and PRC1 as key players in the dynamic regulation of the microtubule cytoskeleton, contributing to a form of cytoskeletal “memory”, where mechanical stress-induced microtubule patterns are reinforced and likely maintained over time. Since both MAP65-1 and PRC1 have conserved functions in cell division, it also remains open whether the nucleation mechanism described here contributes to spindle or phragmoplast formation and has further implications in cell proliferation that were previously unknown. Thus, both proteins have the potential to actively contribute to tissue morphogenesis across the eukaryotic kingdom – particularly in plants, where growth anisotropy mainly relies on the cortical microtubule-cellulose deposition nexus.

## Methods

### MAP65-1, PRC1 and KIF11 purification

Plasmids encoding His-MAP65-1-His (referred to as MAP65-1) and GFP-MAP65-1-His (referred to as GFP-MAP65-1) with *MAP65-1* sequences from *Arabidopsis thaliana* were previously generated^24^. The recombinant proteins were purified from Rosetta 2(DE3) *E. coli* cells. Bacteria were grown at 37 °C to an OD of 0.5 followed by transfer to 20 °C for one hour. Protein expression continued at 20 °C overnight with 0.5 mM IPTG. The next day, cells were collected by centrifugation and frozen. Cell pellets were resuspended in lysis buffer containing 50 mM Sodium Phosphate Buffer, pH 7.9, 200 mM NaCl, 20 mM imidazole, 0.5% Triton X-100, 0.5 mM DTT and a protease inhibitor cocktail (cOmplete, EDTA-free). Samples were then sonicated in a beaker on ice (Bandelin Sonopuls, 9X 20 seconds on/off, 40% duty cycle). The lysate was then centrifuged for 30 min at 16,000 rpm at 4 °C and applied to a column containing Ni Sepharose High Performance beads (Cytiva Life Sciences). The column was washed with a buffer containing 50 mM Sodium Phosphate Buffer, pH 7.9, 100 mM NaCl, 30 mM imidazole and 0.5 mM DTT. The proteins were eluted with 500 mM imidazole and dialyzed overnight in a buffer containing 50 mM Sodium Phosphate Buffer, pH 7.9, 100 mM NaCl and 0.5 mM DTT. Further purification followed by loading the protein on a gel filtration column (HiLoad 16/600 Superdex 200 pg, Cytiva Life Sciences) connected to the NGC Chromatography system (Bio-Rad) in 50 mM Sodium Phosphate Buffer, pH 7.9, 100 mM NaCl and 0.5 mM DTT. Proteins were concentrated using Amicon Ultra-15 Centrifugal Filters (Merck).

Human PRC1 was expressed for 96 h in SF9 cells from a pOCC7 plasmid encoding PRC1 labeled with a His6 tag and a 3C precision cleavage site. For purification, cell pellets were thawed on ice and resuspended in purification buffer (50 mM NaH2PO4, 500 mM NaCl, 2 mM MgCl2, 1 mM DTT, 0.1% Tween20, pH 7.8) with protease inhibitor. The lysate was cleared with an ultracentrifuge spin with 40,000 rpm for 1 h at 4 °C. The supernatant was filtered through a 0.45 μm filter and loaded on a 1 mL HiTrap column with a superloop. The column was washed with IMAC wash buffer (purification buffer with 20 mM imidazole) and the protein was eluted with IMAC elution buffer (purification buffer with 300 mM imidazole) with an elution gradient. Protein-containing fractions were pooled and concentrated with Amicon filters (cutoff 100 kDa). 3C protease was added (1:150, v/v) and the His6 tag was cleaved overnight at 4°C. The protein solution was diluted 6-fold to reduce the imidazole concentration and passed over the HiTrap column again. The protease remained bound to the column with its His6 tag. The flow through was concentrated to 0.5 mL, cleared at 17,000 g for 10 min and gel-filtered over a Superose6 column with purification buffer. 10% glycerol was added, and the protein was flash frozen in liquid nitrogen and stored at -80°C. Human KIF11 was expressed and purified as previously described^49^.

### Tubulin purification and labeling

Fresh bovine brains were used as the source of brain tubulin, which was purified by three cycles of temperature-dependent polymerization and depolymerization in Brinkley Buffer 80 (BRB80 buffer; composed of 80 mM PIPES, pH 6.8, 1 mM EGTA and 1 mM MgCl_2_ supplemented with 1 mM GTP). We obtained MAP-free tubulin by using low (32.6% glycerol, 1.5 mM ATP, 0.5 mM GTP, 3 mM MgCl_2_) and high salt buffers (High Molarity PIPES buffer; 1 M PIPES, pH 6.9, adjusted with KOH, 10 mM MgCl_2_, 20 mM EGTA) and cation-exchange chromatography (EMD SO, 650 M, Merck) in 50 mM PIPES, pH 6.8, 1 mM MgCl_2_ and 1 mM EGTA.

For the labeling (with ATTO-488, ATTO-565, Alexa-Fluor-647 or biotin), microtubules were polymerized with purified brain tubulin at 37 °C for 30 min and layered onto cushions of 0.1 M NaHEPES, pH 8.6, 1 mM MgCl_2_, 1 mM EGTA, 60% glycerol, followed by sedimentation by ultracentrifugation at 37 °C. Microtubules were then resuspended in 0.1 M NaHEPES, pH 8.6, 1 mM MgCl_2_, 1 mM EGTA, 40% glycerol and labeled by adding 1:10 volume 100 mM NHS-ATTO (ATTO Tec) or NHS-LC-LC-Biotin (EZ-link, Thermo) for 10 min at 37 °C. Two volumes of 2X BRB80 with 100 mM potassium glutamate and 40% glycerol were used to stop the labeling reaction, followed by microtubule sedimentation onto cushions of BRB80 supplemented with 60% glycerol. Finally, microtubules were resuspended in BRB80, and a last cycle of polymerization and depolymerization was performed before storage.

### Coverslip treatment

Coverslips were cleaned by successive treatment with the following solutions: 30 min acetone and 15 min 96% ethanol followed by two washes with ultrapure water, then 2 hours in Hellmanex III (2% in water) followed by two washes with ultrapure water. Coverslips were then airdried and treated with UV for 25 min. Next, coverslips were incubated for 3 days in a solution containing a 1:9 mix of triethoxysilane-PEG-biotin and triethoxysilane-PEG (30 kDa, Creative PEGWorks) at 1 mg/ml in 96% ethanol and 0.1% HCl with gentle agitation at room temperature. Coverslips were then rinsed once in absolute ethanol and twice in ultrapure water, airdried and stored at 4 °C.

### Microtubule growth, capping and nucleation dynamics

Microtubules seeds were polymerized in a total volume of 100 μl with 6 μM tubulin (labeled with 30% ATTO-565 or Alexa Fluor 647 and 70% biotinylated tubulin) in 1X BRB80 supplemented with 0.5 mM GMPCPP at 37 °C for 1 hour. 2 μl of 50 μM taxol (Merck) were then added followed by incubation at room temperature for 30 min and ultracentrifugation at 156,000 xg at 25 °C for 10 min. Seeds were then resuspended in 1X BRB80 supplemented with 0.5 mM GMPCPP and 1 μM taxol.

A flow cell chamber with a volume of approximately 20 μl was built using double-sided adhesive tape and a glass coverslip functionalized and passivated as mentioned above. The top and bottom pieces were cut into the desired sizes using a diamond engraving pen. Flow chambers were first perfused with 50 μg/ml Neutravidin (Fisher Scientific) in 1X BRB80 for 1 min, followed by passivation with 0.1 mg/ml PLL-g-PEG (PII 20 K-G35-PEG2K, Jenkam Technology) in 10 mM Na-HEPES, pH 7.4, for 1 min and washed with 1X BRB80. Microtubules seeds were then flushed into the chamber. Non-attached seeds were washed out by using 1X BRB80 supplemented with 1 mg/ml casein (BRB80/casein).

Microtubule polymerization with seeds as template was achieved with a mix containing 11 μM tubulin (10 to 20% labeled with ATTO-565 or Alexa Fluor 647) in 0.7X BRB80 and 0.38X MAP buffer (500 mM Phosphate buffer, 1 mM KCl, 10 mM DTT, pH 7.9) supplemented with 1 mM GTP, an oxygen scavenger cocktail (22 mM DTT, 1.2 mg/ml glucose, 8 μg/ml catalase and 40 μg/ml glucose oxidase), 1 mg/ml casein and 0.033% methyl cellulose (1,500 cP, Sigma) at 37 °C. Microtubules were capped by substituting GTP with 0.5 mM GMPCPP (Jena Bioscience) and using 3 µM 100% labeled tubulin (with ATTO-565 or Alexa Fluor 647) at 37 °C. Capping microtubules extends their lifetime for long-term observation (for instance up to 30 min) and overcomes dynamic instability *in vitro*. To observe microtubule nucleation in the presence of PRC1 or MAP65-1, the same buffer as in microtubule polymerization was used, supplemented with 4 µM tubulin 100% labeled with ATTO-488 and the corresponding protein at the desired concentration (the protein stock was diluted in 1X BRB80). Microtubule nucleation dynamics in the presence of PRC1/MAP65-1 was observed for 30 minutes, unless stated otherwise.

For the experiments varying the amount of microtubule defects: for fast growth conditions, 11 µM tubulin was used for polymerization; for slow growth conditions, 5.5 µM tubulin was used.

To observe stabilized microtubules nucleated by varying MAP65-1 concentrations, microtubule incubation with 100% ATTO-488-labeled tubulin and MAP65-1 (in the same buffer as for microtubule polymerization) proceeded for 20 min, followed by a wash buffer (with the same composition as the microtubule polymerization buffer, but without free tubulin and with 20 µM taxol).

### Nucleated microtubule transport by KIF11

Microtubules were polymerized in microcentrifuge tubes as described below with 10% labeled ATTO-565 and with biotin on their caps and seeds. After washing away taxol, microtubule nucleation proceeded for 30 min in the presence of 100 nM MAP65-1 and 4 µM 100% ATTO-488-labeled tubulin followed by flushing of a kinesin buffer containing 10 nM KIF11, 2 µM of tubulin 100% labeled with Alexa Fluor 647 and 0.7X BRB80 supplemented with 1 mM GTP, 1 mM ATP, an oxygen scavanger cocktail (22 mM DTT, 1.2 mg/ml glucose, 8 μg/ml catalase and 40 μg/ml glucose oxidase), 1 mg/ml casein and 0.033% methyl cellulose (1,500 cP, Sigma). Microtubules were observed for 20 min.

### Microtubule polymerization for bending and taxol treatment

Microtubules were first polymerized in microcentrifuge tubes by using 11 µM tubulin (labeled with 10% ATTO-565 or Alexa Fluor 647) in 200 µl of a buffer containing 1.2X BRB80, 0.6X MAP buffer, 0.5 mM GTP and previously polymerized seeds for 40 min at 37 °C. 5 µl of 30 µM taxol were then added, following by centrifugation for 30 min at 15,000 rpm at room temperature. Microtubules were then resuspended in capping mix containing 0.5 µM tubulin (labeled with 60% biotin and 40% ATTO-565 or Alexa Fluor 647) in 1.2X BRB80, 0.6X MAP buffer, 0.5 mM GMPCPP and 10 µM taxol. Stepwise capping of microtubules was achieved by adding 0.5 µM tubulin at a time followed by incubation for 15 min at 37 °C for a total of ten times. Microtubules were diluted 1:200 in BRB80/taxol (1X BRB80, 10 µM taxol) until usage. The same conditions were used for microtubules referred to as taxol pre-treated, which were incubated overnight at room temperature in BRB80/taxol.

Microtubules were flushed into passivated and functionalized chambers as described above, followed by successive addition of 100 µl BRB80/taxol from both ends of the flow chamber in an alternate fashion to cause microtubule bending due to fluid flow, since microtubules could attach via the seed and the cap to the coverslips. Taxol was washed away by using BRB80/casein. To observe GFP-MAP65-1 binding to bent microtubules, MAP65-1 was added with 15% GFP-MAP65-1 and 85% MAP65-1 in the presence of 2 µM non-fluorescent tubulin. To observe microtubule nucleation on bent microtubules, MAP65-1-mediated nucleation proceeded for 20 min followed by stabilization with a wash buffer with no tubulin and 10 µM taxol.

### Microtubule nucleation using annealed microtubule population

Microtubules were polymerized in a buffer containing 1X BRB80, 1.25 mM GMPCPP, 1.25 mM MgCl_2_ and 2.5 µM tubulin (labeled with 20% ATTO-565 or Alexa Fluor 647 and 20% biotin) for 5 hours at 28 °C, followed by addition of 120 µl 1X BRB80 and centrifugation at 13,000 rpm for 15 min. Microtubules were then resuspended in 150 µl of 1X BRB80 and mixed in equal amounts, followed by incubation at 30 °C overnight to allow for annealing to happen. Microtubules were then flushed into flow chambers and MAP65-1-mediated nucleation proceeded by 20 min followed by stabilization with a wash buffer with no tubulin and 10 µM taxol.

### Microtubule nucleation using a population of microtubules with high and low defect regime

For the high defect regime, microtubules were polymerized in 10 µl of a buffer containing 1X BRB80, 2 mM GMPCPP, 0.1 mM MgCl and 20 µM tubulin (labeled with 20% Alexa Fluor 647 and 20% biotin) for 30 min at 37 °C, followed by addition of 190 µl of 1X BRB80 and centrifugation at 156,000 xg for 15 min. Microtubules were then resuspended in 150 µl of 1X BRB80. For the low defect regime, microtubules were polymerized in 80 µl of a buffer containing 1X BRB80, 1.25 mM GMPCPP, 1.25 mM MgCl_2_ and 2.5 µM tubulin (labeled with 20% ATTO-565 and 20% biotin) for 5 hours at 28 °C, followed by addition of 120 µl of 1X BRB80 and centrifugation at 13,000 rpm for 15 min. Microtubules were then resuspended in 150 µl of 1X BRB80. Microtubules were then flushed in sequentially and MAP65-1-mediated nucleation proceeded by 20 min followed by stabilization with a wash buffer with no tubulin and 10 µM taxol.

### Plant growth conditions

*Arabidopsis thaliana* seeds were surface-sterilized by treatment with a solution containing 2% bleach and 0.05% Triton X-100 for 5 min followed by three washes with sterile distilled water and resuspension in 0.05% agarose. Seeds were sown on ½ MS medium (basal salt mixture, Duchefa Biochemie) supplemented with 0.5% sucrose and 0.8% plant agar (Duchefa Biochemie). Plates containing seeds were stratified at 4 °C for 2 to 3 days in the dark. Plants were grown in a 16-hour/21 °C light and 8-hour/18 °C dark regime with 60% humidity.

### Protoplast isolation and observation

Roots from seedlings grown for eleven days were dissected and inserted in an enzyme solution composed of solution A (2 mM CaCl_2_, 2 mM MgCl_2_,10 mM MES, pH 5.5 adjusted with KOH, and 600 mOsmol/l Mannitol) and the following enzymes: 17 mg/ml Cellulysine (Calbiochem), 17 mg/ml Cellulase RS (Duchefa Biochemie) and 0.4 mg/ml Pectyolase from *Aspergillus japonicus* (Sigma-Aldrich). Roots were digested for two to four hours at room temperature on a rotating stage at 15 rpm. Next, the solution was filtered through a 70 µm filter and the filter was washed with 1 ml of solution A. The filtered protoplasts were then centrifuged for 4 min at 1,000 rpm. The supernatant was removed and 1 ml of solution A was added, followed by gentle flicking. The protoplasts were centrifuged again for 4 min at 1,000 rpm, followed by supernatant removal and resuspension in 200 µl of solution A. The protoplasts were then applied on microwells and allowed to sediment for 10 min. 3 ml of solution B (the same as solution A, but with 280 mOsmol/l Mannitol) were then added to promote protoplast pressurization. Microscopic observation started 2 hours after treatment with solution B and continued for another 2 hours. The 12 X 40 µm microwells were produced as previously described^25^. We selected protoplasts that had an aspect ratio of at least 1.1 (major axis divided by minor axis) to make sure that only enclosed protoplasts were taken into account for our analysis. We included protoplasts with an aspect ratio range of 1.28 to 1.12.

### Imaging conditions and image analysis

To observe plant cells, a point scanning confocal microscope (Zeiss LSM 900) with Airyscan 2 and Axiocam 705 camera was used. *In vitro* microtubules were either visualized with the Zeiss LSM 900 with a stage that was kept at 37 °C through a cage incubator (PECON), or an objective-based orbital TIRF microscope (Nikon Eclipse Ti2, modified by Visitron Systems) and EMCCD camera (Andor iXon Life) at minimal laser intensity with a stage kept at 37 °C through a warm stage controller (OkoLabs). For the Zeiss LSM 900 microscope, the ZenBlue software version 3.2 was used. For the TIRF microscope, the VisiView software version 6.0 was used.

To observe microtubule nucleation dynamics in the presence of PRC1/MAP65-1, images were taken every 4 s for a total of 30 minutes. Nucleation dynamics were observed through kymographs. The distance between nucleation events was measured manually using the segmented line on Fiji.

To distinguish between incorporation and nucleation events in the case of stabilized microtubules, typically 50 images were taken per field of view (FOV) with an interval of 0.5 to 1 s. Images were processed for background subtraction and smoothing. Line scans of the green fluorescence intensity were drawn along maximum intensity projections of individual microtubules. The elongation of microtubule ends was used as a reference for a full microtubule for every FOV – an event that surpassed the intensity found at the polymerized microtubule ends was scored as a nucleation event if it co-localized with the microtubule lattice.

To quantify the recruitment of free tubulin on the microtubule lattice by MAP65-1, a 1-µm section in length spanning the whole microtubule width was drawn using the polygon tool in Fiji. This section was moved along the microtubule from the beginning to the end. For each individual microtubule, the integrated density measured from each microtubule section was divided by the integrated density in a background region with the same area right next to the microtubule.

To assess whether microtubule nucleation sites overlapped with microtubule annealing sites, we measured the distance between the peak maximum of the nucleation fluorescence signal and the nearest annealing site. If this distance was less than 500 nm, the nucleation site was classified as overlapping with the annealing site. As a control, we selected 55 random 500 nm-long patches along annealed microtubules. A patch was scored as containing a nucleation event if the peak maximum of a nucleation fluorescence signal fell within the 500 nm interval.

### FRAP assay

The GFP-MAP65-1 and free tubulin signals were bleached by using 5 cycles of the 405 laser line at 100% at a speed of 50 ms per pixel. Images were acquired every 0.9 s for 5 min. Bleaching events that resulted in less than 30% of the initial fluorescence intensity were used for the analysis. The fluorescence intensity was normalized to the initial maximum value and plotted over time by using the Stowers Plugins Collection (https://research.stowers.org/imagejplugins) after background subtraction (rolling ball 50 pixels).

### Microtubule curvature and orientation analysis

Microtubules were tracked with the JFilament Fiji plugin (https://imagej.net/plugins/jfilament). Using custom-written python code, the curves of the tracked filaments were first smoothed by parametric Spline interpolation. Then, the menger curvature was computed and the orientation was calculated via finite difference. GFP-MBD and MAP65-1-RFP fluorescence values were normalized for every protoplast against the mean fluorescence intensity (for each corresponding channel) in a circle with a diameter of 2 µm drawn in a region of the cytoplasm with no microtubules present.

### Statistical methods

All statistical analyses were performed with GraphPad Prism. For the spatial frequency quantifications, microtubules for each experimental condition were concatenated in a random order and the distance between two adjacent nucleation events was measured (similarly to what was done previously for the quantification of spatial incorporation frequency^32^).

Data sets were tested for their normality. All of the tested datasets had at least one group that was non-normally distributed, thus non-parametric tests were employed to test if differences were statistically significant (Mann-Whitney and Kruskal-Wallis).

To test if the microtubule nucleation frequency differed in regions containing microtubule annealing sites compared to randomly selected patches of the annealed microtubule population, a Fisher’s exact test was used.

## Data availability

The datasets generated and analyzed in this study are available from the corresponding authors upon request.

## Acknowledgments

MRM and OH were supported by a grant from the European Research Council (ERC, grant agreement number 101019515, “Musix”, awarded to OH). MRM was also supported by a grant from the German Research Foundation (DFG, project number 545084341, awarded to MRM). LS was supported by the DFG grant SFB 1027 and the ERC grant StG 101115795 “CROSSTALK”.

## Author contributions

MRM, LS, OH and SD conceived and guided the project. MRM, LS, OH and SD designed the experiments. MRM and CG performed the experiments. MG and BK purified the tubulin. LM, LN and SD provided KIF11 and PRC1 and related expertise. MRM and CG analyzed the experiments. MRM, LS and OH wrote the manuscript with input from all authors.

---

**Fig. S1, related to Fig. 1:**
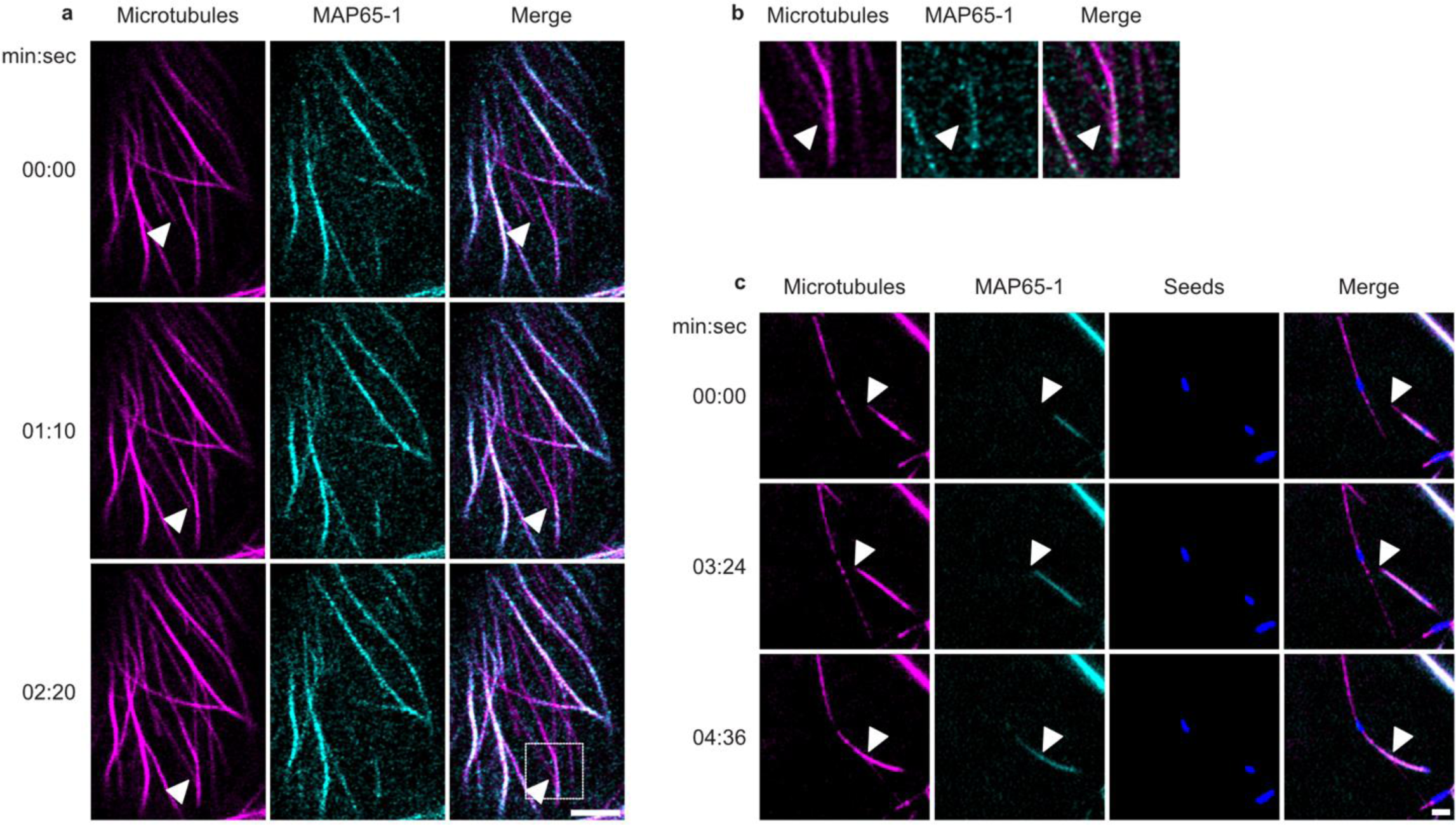
MAP65-1 is a well-known microtubule bundler. **a,** Confocal laser scanning microscopy (CLSM) of epidermal hypocotyl cells from 5-day-old plants co-expressing *p35S::GFP-MBD* and *pMAP65-1::MAP65-1-mCherry*. White arrowhead indicates a microtubule that grows and gets bundled in the last time point (marked with a dashed white square). Scale bar, 5 µm. **b,** Close-up of dashed white square indicated in panel **a**. **c,** *In vitro* microtubule dynamics imaged by total internal reflection fluorescence microscopy (TIRFM) in the presence of 8 μM ATTO-565-labeled free tubulin (magenta) and 50 nM GFP-MAP65-1 (cyan). Stable GMPCPP seeds labeled with Alexa Fluor 647 appear in blue. White arrowhead indicates a microtubule that grows and gets bundled in the last time point. Scale bar, 5 µm.

**Fig. S2, related to Fig. 1:**
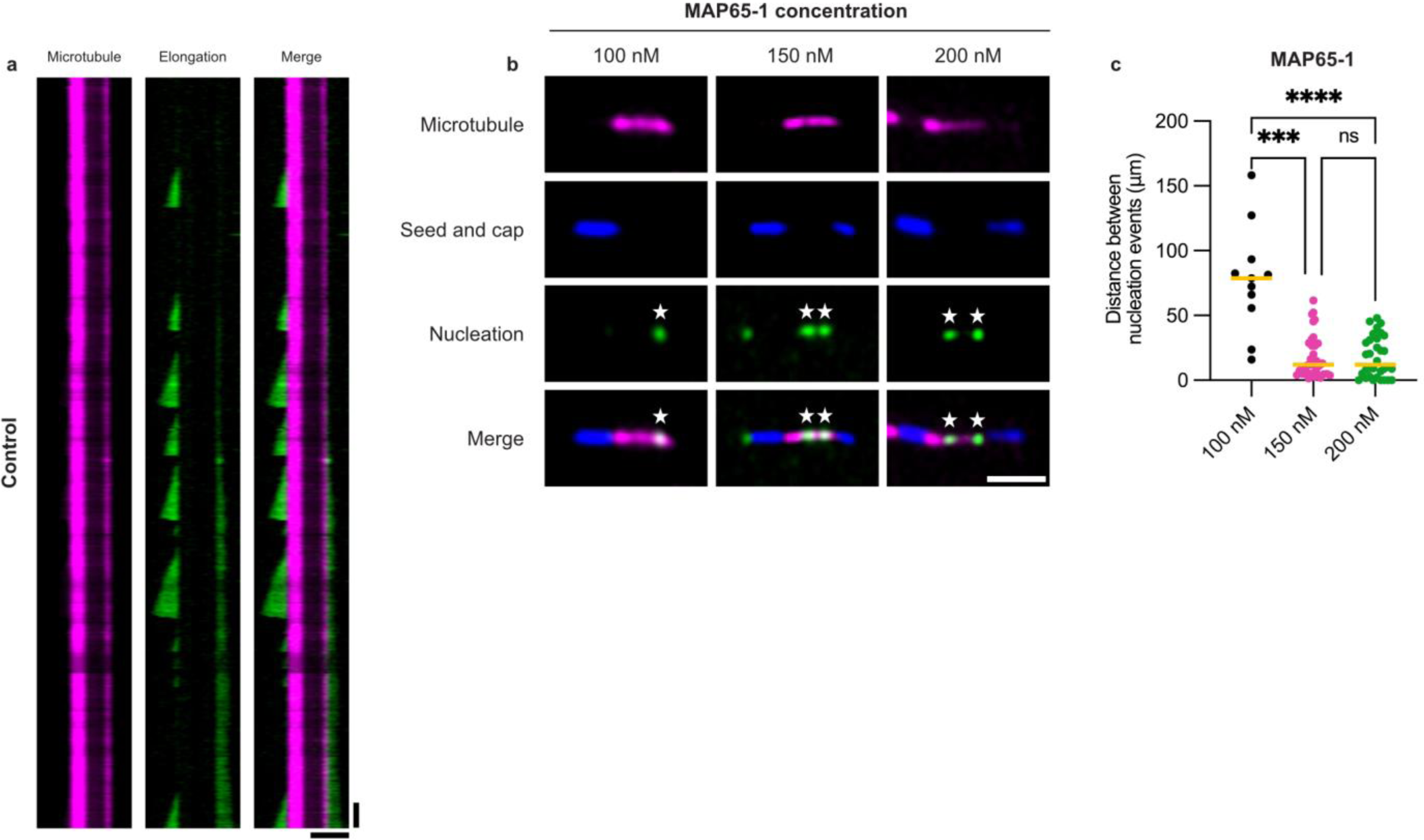
Nucleation does not happen in the control and is MAP65-1 concentration-dependent. **a,** Kymograph of the corresponding control microtubule shown in Fig. 1c (top panel), exhibiting growth from the free ends in the absence of MAP65-1/PRC1. Free tubulin labeled with ATTO 488 (green) and microtubule labeled with ATTO 565 (magenta). Scale bars, 5 µm (horizontal) and 1 min (vertical). **b,** TIRFM images showing microtubule nucleation by MAP65-1 with different concentrations. White stars indicate nucleation events. Scale bar, 5 µm. **c,** Quantification of the distance between nucleation events using different MAP65-1 concentrations. A total of 11 (100 nM), 34 (150 nM) and 32 (200 nM) nucleation events were observed. Bars indicate the median values. Total microtubule lengths of 1009.49 µm, 705.76 µm and 587.54 µm were analyzed respectively, N = 3. Statistics: Kruskal-Wallis test, *** P = 0.0001 and **** P < 0.0001.

**Fig. S3, related to Fig. 1:**
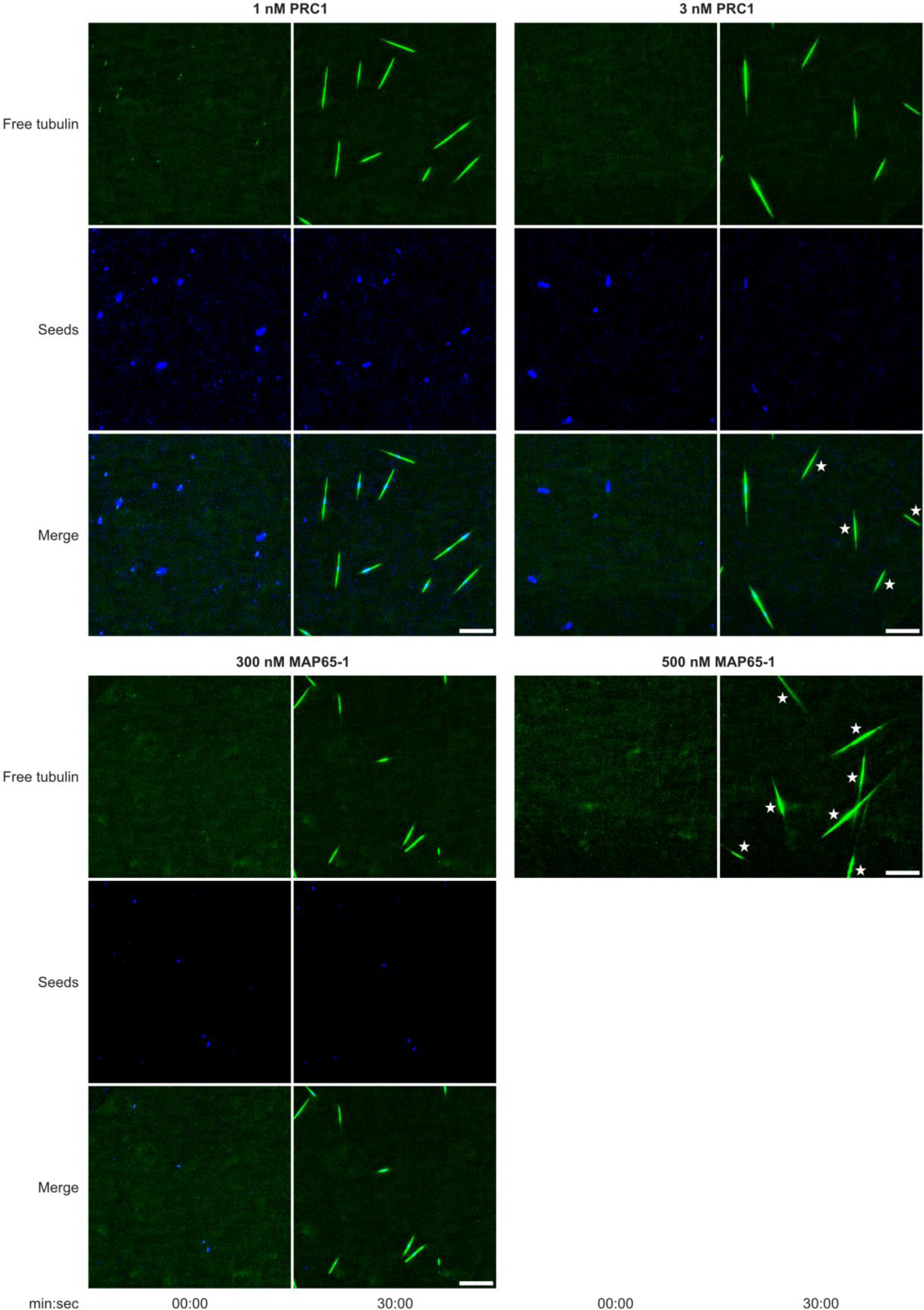
Microtubule nucleation thresholds in solution for MAP65-1 and PRC1. 4 µM ATTO-488-labeled free tubulin was incubated either in the presence of PRC1 or MAP65-1 with the indicated concentrations. For the condition of 500 nM MAP65-1, no seeds are visible. White stars indicate nucleation events in solution. Scale bars, 25 µm.

**Fig. S4, related to Fig. 1:**
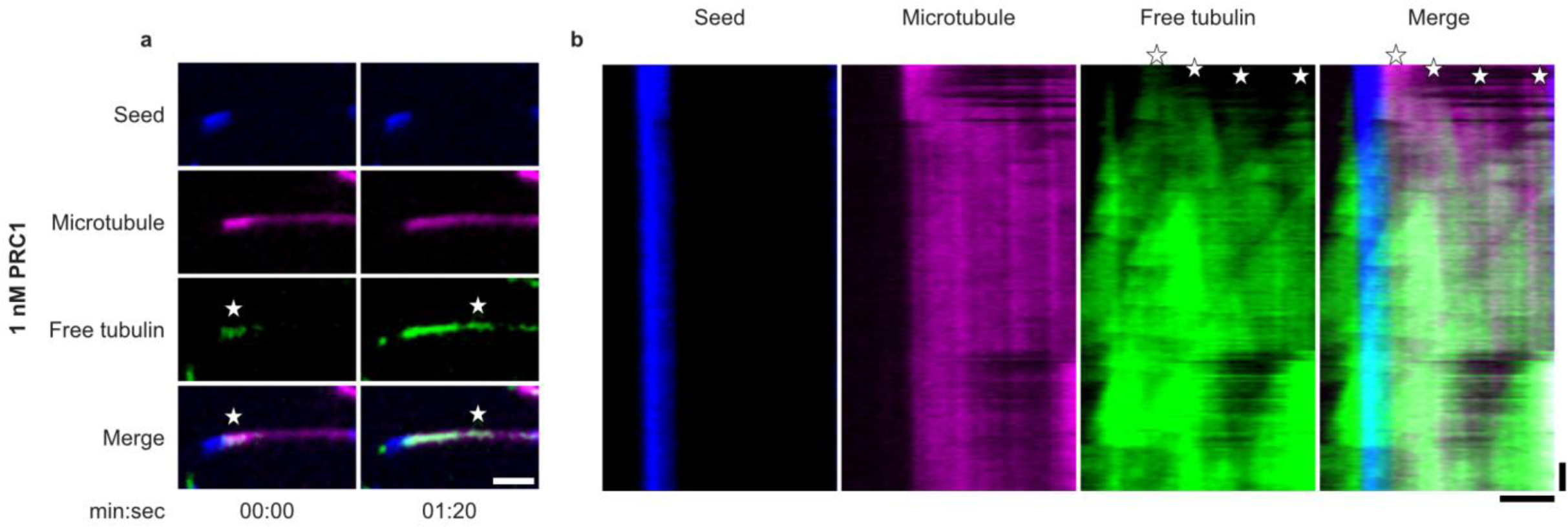
1 nM PRC1 promotes extensive nucleation. **a,** Close-up of microtubule from Fig. 1k showing nucleation events (white stars) along the microtubule lattice in the presence of 4 µM free ATTO-488-labeled tubulin (green) and 1 nM PRC1. Scale bar, 5 µm. **b,** Kymograph of corresponding microtubule shown in **a**. White stars indicate nucleation events. Scale bars, 5 µm (horizontal) and 1 min (vertical).

**Fig. S5, related to Fig. 2:**
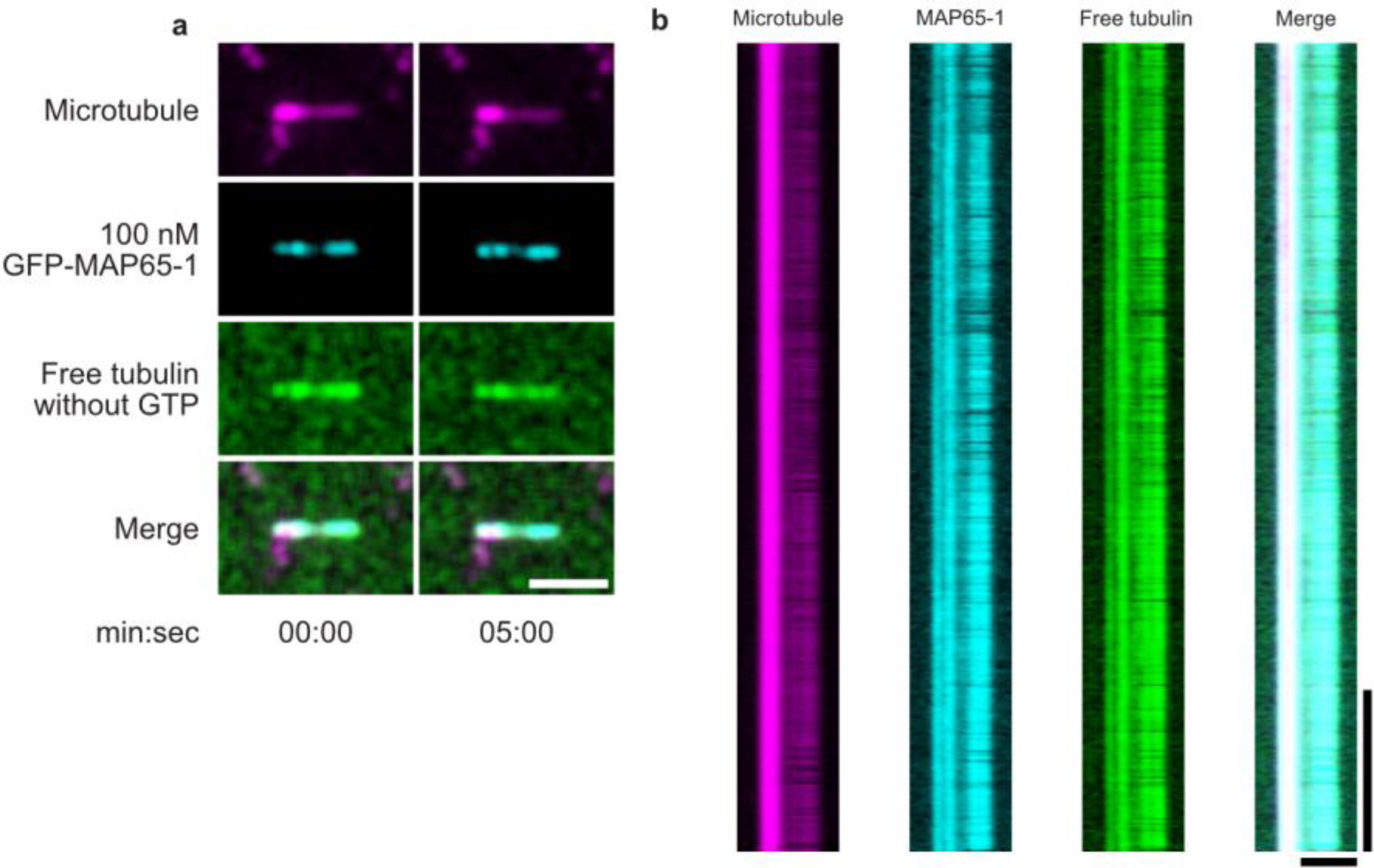
In the absence of GTP, no growth happens from microtubule ends. **a,** Microtubule incubated with 100 nM GFP-MAP65-1 and 4 µM free ATTO-565-labeled tubulin in the absence of GTP imaged by TIRFM. Scale bar, 5 µm. **b,** Kymograph of the corresponding microtubule shown in **a**. Scale bars, 5 µm (horizontal) and 1 min (vertical).

**Fig. S6, related to Fig. 4:**
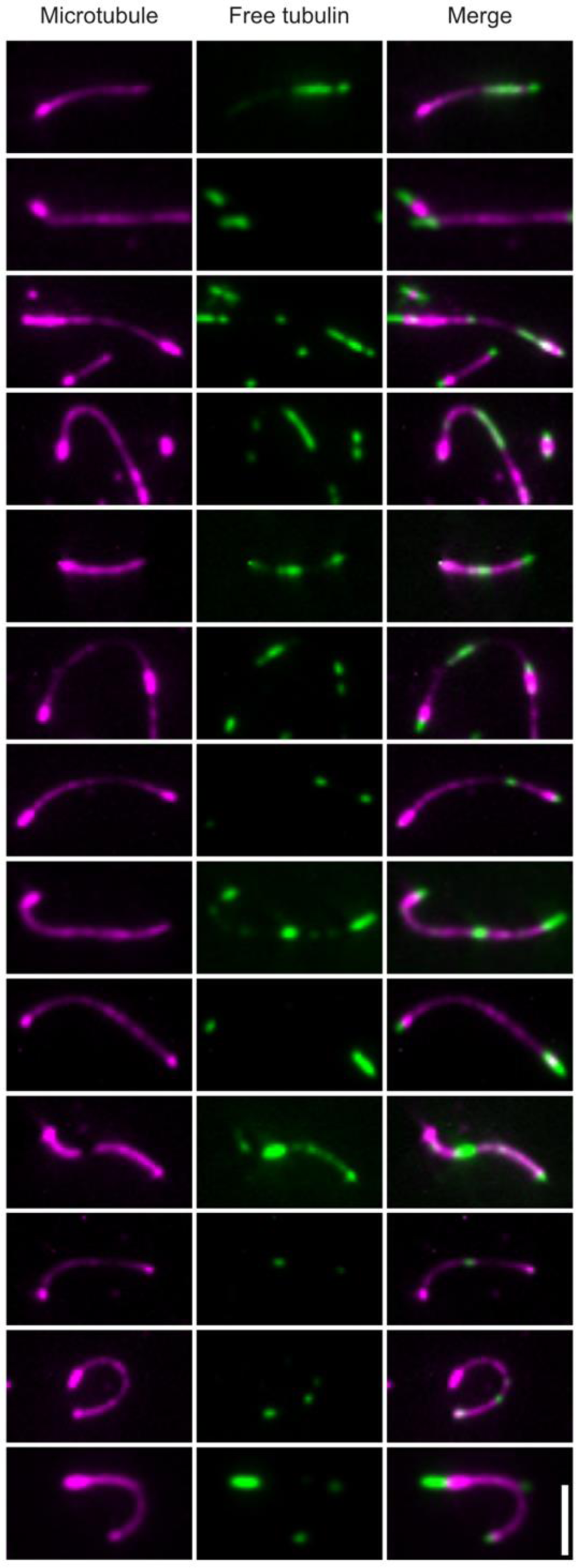
Nucleation events do not always happen at the highest curvature regions. Examples of site of microtubule nucleation on bent microtubules, showing the stochastic nature of this event. The nucleation does not always happen at the site of highest microtubule curvature. Scale bar, 5 µm.

**Fig. S7, related to Fig. 4:**
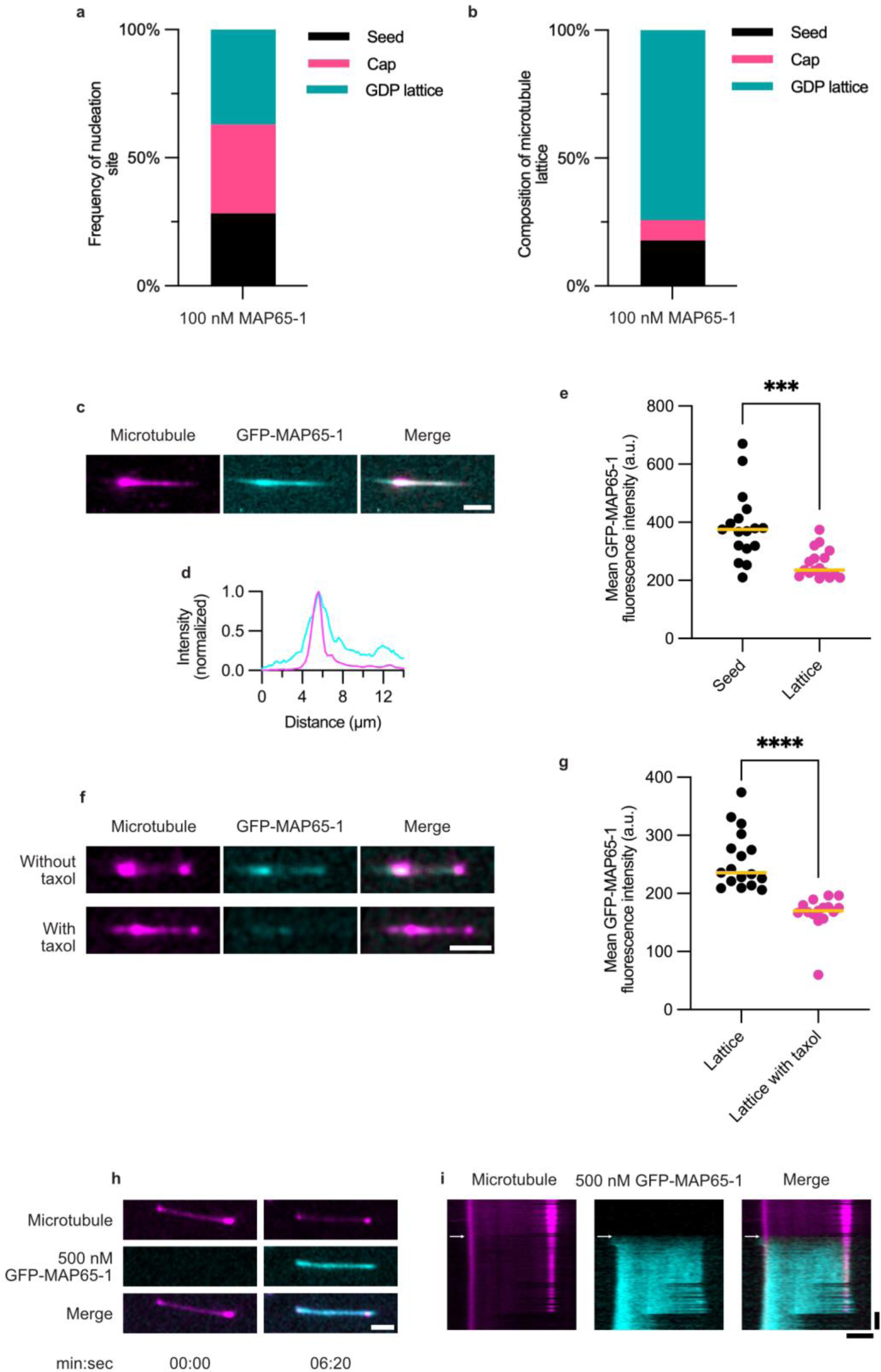
MAP65-1 does not specifically recognize an expanded lattice. **a,** Quantification of the frequency of the nucleation origin site in microtubules grown with 11 µM of 10% ATTO-565-labeled tubulin and incubated with 100 nM MAP65-1 and 4 µM ATTO-488-labeled tubulin as shown in Fig. 1a. n = 46 nucleation events, N = 4. **b,** Average composition of the microtubule lattice in length from the microtubules that were used for the quantification in **a.** Total microtubule length analyzed (GMPCPP seed, GMPCPP cap, and GDP lattice) = 578.94 µm. **c,** Microtubule incubated with 100 nM GFP-MAP65-1 and 3 µM black tubulin and observed by TIRFM. Scale bar, 5 µm. **d,** Graph represents line scans along microtubule shown in **c**, with MAP65-1 in cyan and the microtubule in magenta. The peak corresponds to a GMPCPP region of the microtubule, the seed. **e,** Quantification of GFP-MAP65-1 intensity on the seed (GMPCPP) and lattice (GDP) portions of the microtubule, n = 17 microtubules. Bars indicate the median values. Statistics: Mann-Whitney test, *** P = 0.0001. **f,** Microtubules incubated with 100 nM GFP-MAP65-1 and 3 µM black tubulin or with 20 µM taxol observed by TIRFM. Scale bar, 5 µm. **g,** Quantification of GFP-MAP65-1 intensity on the GDP lattice with and without 20 µM taxol, n = 15-17 microtubules. Bars indicate the median values. Statistics: Mann-Whitney test, **** P < 0.0001. **h,** Stills of a microtubule before and after addition of 500 nM GFP-MAP65-1 and 4 µm black tubulin imaged by TIRFM in a microfluidics setup. Scale bar, 5 µm. **i,** Kymograph of the corresponding microtubule shown in **h**, showing no obvious change in microtubule length upon binding of MAP65-1. White arrow indicates moment in which MAP65-1 is flushed in. Scale bars, 5 µm (horizontal) and 1 min (vertical).

**Fig. S8, related to Fig. 5:**
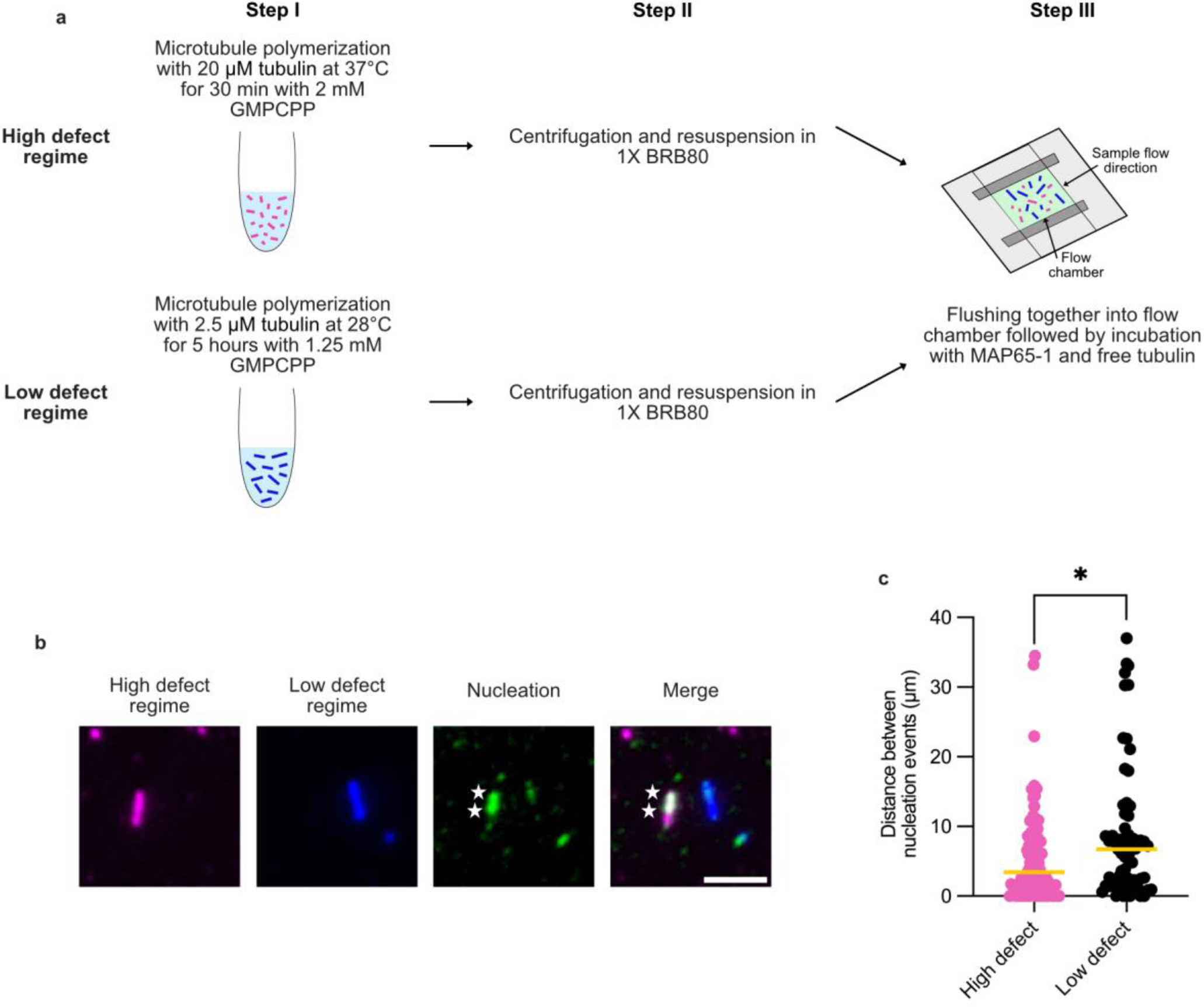
Microtubule nucleation frequency depends on the amount of microtubule defects. **a,** Schematic representation of the experimental setup to obtain microtubules with high and low amounts of defects. Microtubules are either polymerized in a high or low defect regime (step I), labeled respectively with 20% Alexa Fluor 647 and 20% ATTO 565. Next, 1X BRB80 is added and microtubules are centrifuged and resuspended in fresh 1X BRB80 (step II). Microtubules are then sequentially flushed in the flow chamber for incubation with 100 nM MAP65-1 and free tubulin (step III). **b,** Microtubule nucleation mediated by 100 nM MAP65-1 on microtubules polymerized under a high or low defect regime imaged by TIRFM. Scale bar, 5 µm. **c,** Quantification of the distance between nucleation events on microtubules polymerized under a high or low defect regime. A total of 95 (high defect) and 67 (low defect) nucleation events were observed. Bars indicate the median values. Total microtubule lengths of 545.41 µm and 626.598 µm were analyzed respectively, N = 3. Statistics: Mann-Whitney test, * P = 0.0433.

